# Local mating competition, but not climate, drives male reproductive success across a latitudinal gradient in a nest-brooding marine fish

**DOI:** 10.64898/2026.02.10.705067

**Authors:** Ivain Martinossi-Allibert, Yimen Gerardo Araya Ajoy, Sebastian Wacker, Trond Amundsen

## Abstract

Understanding ecological drivers of reproductive success is crucial to predict whether natural populations can cope with the pace of anthropogenically driven environmental change. In marine ecosystems, this knowledge is difficult to acquire due to the lack of tractable field systems. Here, we took advantage of the nest-brooding behavior of the two-spotted goby *Pomatoschistus flavescens*, an important planktivorous fish in Scandinavian coastal ecosystems, to study its reproduction across the steep climatic gradient of its natural range. We deployed 360 artificial nests in the field, covering six populations during the breeding season of 2022. We found that climate explained differences among populations in the phenotypes of nest-holding males, and in the impact of both marine growth and parental cannibalism on the broods. In addition, climate affected egg density and diameter. Despite these ecological effects, and although populations differed in average male reproductive success, reproductive success was not influenced by climate. Instead, it was largely determined by competition occurring at the local scale, in particular through the acquisition of high-quality nests, which was itself affected by the relative size of males within the local pool.

We propose that the frequency-dependent nature of mating competition may buffer reproductive success against climatic influence in *P. flavescens*, and discuss the potential generality of such mechanisms and implications for population resilience.

## Introduction

In a world that is changing rapidly under anthropogenic pressure (Wiens 2016), biologists are tasked with predicting impact on ecosystems at various scales, as well as assessing the resilience of wild populations (e.g, Pinsky et al. 2019). In this context, reproductive dynamics matter because of their role in population persistence, but their sensitivity to rapidly changing environments is not well understood (Candolin and Heuschele 2008, Pilakouta and Ålund 2021). For instance, a breakdown of mating communication due to environmental disturbances (e.g., Candolin et al. 2007) could lead to rapid demographic decline. Conversely, efficient sexual selection can, under certain circumstances, accelerate adaptation, and thus contribute to evolutionary rescue (Cally et al. 2019, but see Martinossi-Allibert et al. 2019b). Furthermore, because mating is inherently a competitive activity, it is not straightforward to predict how ecological effects on individual traits may extrapolate into population-level dynamics, and ultimately affect population growth and likelihood of evolutionary rescue (Svensson and Connallon 2019). For this reason, it is necessary to study ecological effects on mating interactions and reproductive success in population context, a task that is particularly difficult in marine ecosystems. To address this challenge, we take advantage of the nest-brooding behavior commonly associated with male-only parental care in teleost fish (Blumer 1982). Nest-brooding makes the study of reproductive dynamics in the field tractable, and nest-associated traits can be valuable indicators for environmental sensitivity (Mainwaring et al. 2017).

Anthropogenic impact on marine ecosystems includes changes in water temperature and pH (Cheng et al. 2019, Doney 2010, Miller et al. 2013), noise (Peng et al. 2015) and turbidity (Lunt and Smee 2020). Several important avenues of research have already been opened, aimed at understanding the impact of these changes on fish reproduction. Physiologists are particularly interested in the effects of temperature, which affects phenology and sex determination pathways (e.g., Lema et al. 2024). By observing behavior in the laboratory, researchers have also demonstrated the sensitivity of sexual signals to environmental perturbations such as temperature (Albouy et al. 2023), noise (de Jong et al. 2018) or turbidity (Järvenpää and Lindström 2004, Sundin et al. 2017, Järvenpää et al. 2019). The effects of climate change on commercially exploited pelagic stocks are also well documented, but often restricted to population growth, phenology, geographical distribution, and body size (McKenzie and Geffroy 2021). Important aspects of the reproductive dynamics of marine fish thus remain understudied. In particular, field studies on non-commercial species are lacking, although such species can have important roles in their ecosystems (but see: Laur et al. 2014, Monteiro et al. 2017, Martinossi-Allibert et al. 2025a).

Assessing temperature effects on reproductive dynamics is particularly relevant for high latitude marine ecosystems of the northern hemisphere, where ocean temperature is predicted to rise the fastest (Ruela et al. 2020). We leveraged our experience with the two-spotted goby *Pomatoschistus flavescens* system to collect data on nest guarding males and their broods in natural populations spread across a steep climatic gradient in the North-East Atlantic. In the spring and summer of 2022, we conducted a large-scale field survey involving six study populations, along the Swedish and Norwegian coastlines, from Lat. 58°N to Lat. 68°N. Each population was represented by three study sites (18 sites in total), and each site was sampled twice, first at the peak of breeding season (June), and then late in the breeding season (July). We used artificial nests of two different sizes, allowing variation in nest quality along this one controlled dimension.

In order to understand the effect of climate on the reproductive strategy of *P. flavescens*, and to disentangle the drivers of individual reproductive success, we used the field data to inform a path diagram that describes the relationships between thermal environment, male traits (size, condition), nest quality, nest status (overgrowth, parental cannibalism), and reproductive success.

## Methods

### 1. Study species and nesting behavior

The two-spotted goby *Pomatoschistus flavescens* is a semi-pelagic fish, ubiquitous in the coastal ecosystems of the North-East Atlantic (Collins 1981), and typically reaching between 4 to 6 cm in adult size (Amundsen 2018). Individuals breed near the shore during spring and summer (April to August in Scandinavia, Skolbekken et al. 2001). Both sexes show colorful ornaments and perform elaborate courtship displays (Forsgren et al. 2004, Wacker et al. 2013). Along most of its range, *P. flavescens* is an annual species, with both sexes able to breed repeatedly during a single season. However, some individuals may reproduce two consecutive years (Johnsen 1945), and in populations to the north of the species range (above Lat. 63°N) postponement of reproduction to the second year of life may be common (Martinossi-Allibert et al. 2025a).

Parental care is performed exclusively by males, from fertilisation to hatching of the eggs (Skolbekken & Utne-Palm 2001). Males hold a nest in a natural cavity such as an empty mussel shell or a fold in a kelp leaf (Mobley et al. 2009, Amundsen 2018).

Each nest holder receives eggs from multiple females, but in most cases all eggs in the same nest develop synchronously and hatch within 24h (Mobley et al. 2009, de Jong 2011). We refer to the eggs provided by one female as a clutch, and the total mass of eggs in a nest, usually composed of several clutches, as a brood, following de Jong (2011). Females visit several nest-holding males before spawning (Myhre et al. 2012), but likely lay their entire clutch in one nest (Skolbekken & Utne-Palm 2001). Interfering behaviours from other males can occur: sneaker males may attempt to fertilise part of the clutch (Utne-Palm et al. 2015), and nest take-overs are also observed (de Jong 2011).

After fertilisation of the eggs, the attending male performs parental care until hatching, by cleaning and oxygenating the eggs and protecting them against predators (Skolbekken & Utne-Palm 2001, Bjelvenmark and Forsgren 2003). The development time of the eggs is temperature dependent, with laboratory estimates giving 15 days at 13°C and 10 days at 18°C for individuals of the Swedish populations of Kristineberg (Bjelvenmark and Forsgren 2003).

### 2. Sampling design

We included six study populations, from the west coast of Sweden (Kristineberg population, Lat. 58°N) to the north of the Lofoten in Norway (Ringstad population, Lat. 68°N, see Figure 1.a and Supplementary File F1). The populations were chosen to be spread evenly across the latitudinal gradient, while giving priority to sites where P. flavescens had been previously studied. For each population, three replicate nest lines were placed in different sites, located within a few kilometers of each other. Study sites were close to the shore, with 1-2m of water depths at low tide, and within kelp and seaweed habitat, which is suitable for the breeding of *P. flavescens*. Each nest line was constituted of 20 artificial nests, alternating large and small nests, and placed about 2 meters from each other, within the constraints imposed by the local underwater landscape. In total, 360 artificial nests were placed in the field, 60 per study population (Figure 1). Each nest was sampled twice, first during peak breeding season and then during late breeding season.

**Figure 1.**
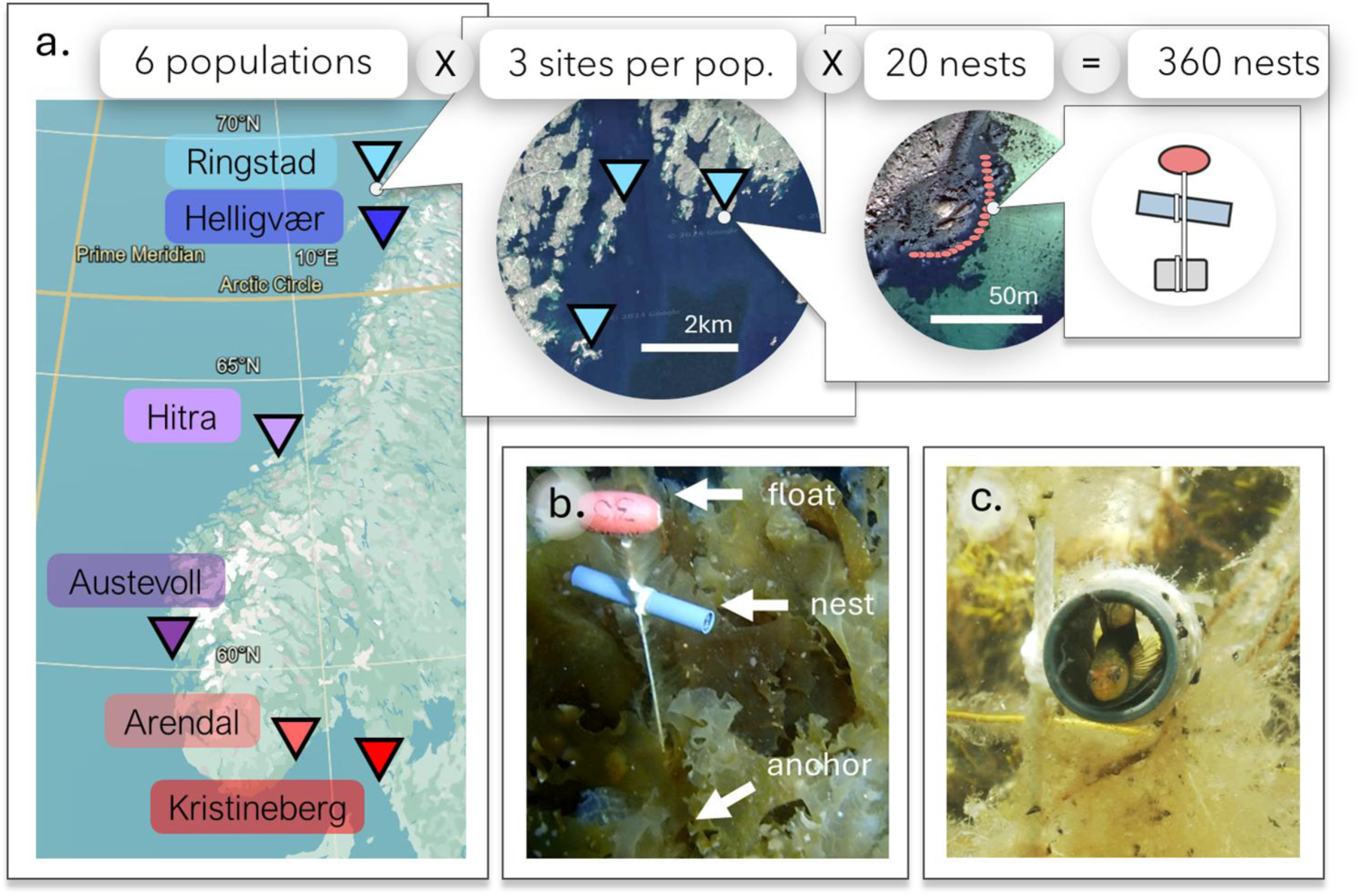
Sampling design and location of the six study populations (a), artificial nest in the field (b) and nest-guarding male (c). In (a), the location of the six study populations is shown (background map from Google Earth 2024). Each population comprised three replicate nest lines (sites) of 20 artificial nests each, located within a few kilometers of each other, representing a total of 360 nests. See supplementary file F1 for the detailed location of each nest line. In (b), an artificial nest photographed in the field (photograph by Trond Amundsen). In (c), a male oxygenating his clutch in an artificial nest (photograph by Ivain Martinossi-Allibert).

### 3. Thermal environment

To represent how the latitudinal variation in climate affected the thermal energy available for growth and reproduction in each of the six *P. flavescens* populations, we used a simple index created by Martinossi-Allibert et al. (2025a). Briefly, this index estimates the area under the water temperature curve for each population in the breeding habitat over a fixed period encompassing the breeding season (1st of April to 31st of August). The temperature data was obtained from the Norkyst model (Asplin et al. 2020) on the period 2005-2023, and monthly averages were calculated for each population. A sinusoidal function was fitted to these points for each population, and integrated over the desired period, giving the DDF index, in units of degree days. The sinusoidal fit is shown on Figure 2a and the DDF index for each population in Figure 2b.

**Figure 2.**
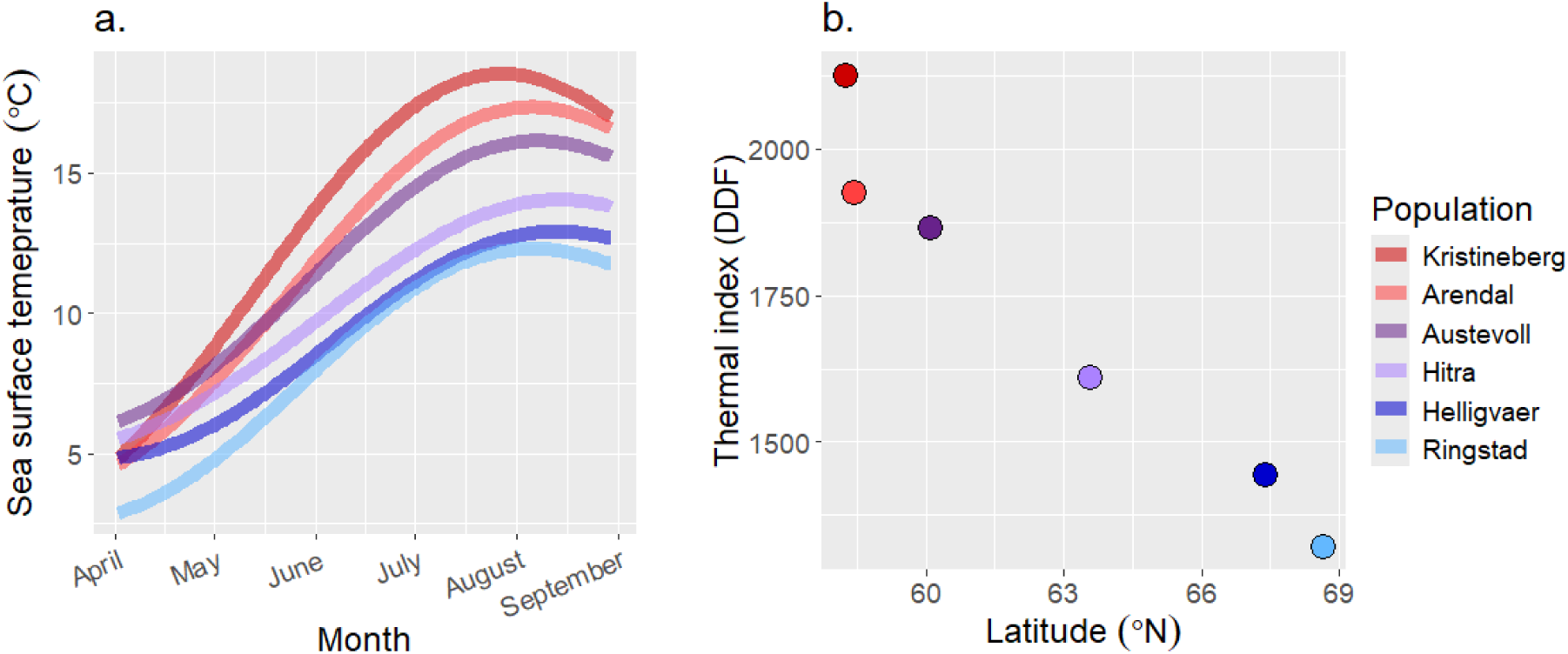
Thermal environment in the six P. flavescens populations: average sea surface temperature (a) and thermal index (b). The sea surface temperature in (a) is a sinusoidal fit to data from the Norkyst model (Asplin et al. 2020) for the six populations averaged over the period 2005-2023. To represent the thermal environment with a simple index, the temperature curves were integrated over a fixed period representing the breeding window, from 1st of April to 31st of August. This gives the Degree Days Fixed (DDF) index shown in (b), see Martinossi-Allibert et al. 2025a for further details.

### 4. Artificial nests

Artificial nests in the field are easily located upon return to a field site, despite the rapid growth of local vegetation. Moreover, they facilitate comparisons across sampling times and latitudes because they present consistent nest quality, while the quality and abundance of natural nests may vary. Artificial nests were composed of a PVC tube, which two-spotted gobies have been found to easily take over as nests in previous experiments (de Jong 2011, Mück et al. 2013, Amundsen 2018). The PVC tubes were lined with a transparent sheet of acetate, which allows for easy extraction and photography of the nest content. The acetate sheets were printed with a grid pattern to facilitate surface estimation. For each study site of 20 nests, we provided 10 small nests (length=80mm, internal diameter=16mm) and 10 large ones (length=175mm, internal diameter=20mm), in order to test the effect of this aspect of nest quality on male reproductive success and male phenotype. Larger nests are assumed to be of higher quality, because they can hold more eggs. The nests were anchored to the bottom by a brick, to which the PVC tube was connected by a line. The line continued above the tube and was attached to a float marked with an ID number and allowing for easy retrieval of the nest (Figure 1.b).

### 5. Collection and measurements of males and nests

The nests were placed in the field in the spring of 2022 between the 24th of April and the 3rd of June. Nest data was then collected twice, first during peak breeding (June) and then during late breeding season (July). Between nest placement and data collection, and between the two rounds of data collection, each study site was left unvisited for about 15 days to allow for the settlement of males (see calendar in Supplementary file F1). Data collection was performed by a snorkeling field worker, assisted by a team on land. The snorkeler started by locating the nests and observing if a guarding male was present (a guarding male in his nest typically peaks out to observe the approaching snorkeler before retreating inside). Males occasionally go on foraging trips during parental care; thus, a nest was recorded as not occupied only after several checks performed over 2-3 hours. To capture a male, the snorkeler used two dip nets on each side of the PVC tube. Males were brought to shore and held in a container of fresh sea water. The nest was then retrieved to shore and the acetate sheet lining the inside of the tube extracted and stored in a container with fresh sea water. After the first sampling, the acetate lining was replaced with a new one and the nest was replaced and left to be colonised by a new male. After the 20 artificial nests of a study site had been checked, the males and nests collected were brought to the field station for measurements. Males were measured for length to the nearest half millimeter using a measuring board, and weighed to the nearest centigram using an electronic scale. The males were then euthanized in excess of MS 222 and preserved in 95% ethanol. The broods extracted from the nests were photographed before being preserved in 95 % ethanol. These brood photographs were later used to estimate male reproductive success, as well as cannibalism and overgrowth indices. During field collection, the occupancy status of the nests was established as (i) having a guarding male or not and (ii) having live eggs or not.

### 6. Male condition

Male condition was estimated as the residuals of a linear regression between the logarithm of weight, as a response variable, and the logarithm of the total body length, as an explanatory variable (following de Jong 2011).

### 7. Picture analysis of nests

Pictures of the content of the nests were analysed after the field collection, in January and February 2025. Examples of typical pictures from the dataset are shown in Supplementary Figure S1. The goal of the picture analysis was to estimate male reproductive success (Egg number), as well as aspects of nest status (brood cannibalism and overgrowth). The detailed image analysis protocol with example pictures and an assessment of measurement repeatability are given in Supplementary File F2, Part 1 and 2.

#### Exclusion of nests

Two main criteria were used to exclude nests (Supplementary File F2, Part 1.1): (i) if eggs were deemed to be dead or if (ii) the brood started hatching during collection. If the eggs were reported as dead on the field form, or if the eggs appeared white and mushy in the picture, the nest was excluded from the analysis. Nests with dead eggs are likely abandoned and subjected to predation, not giving relevant estimates of nest traits. When the egg development stage was very advanced, eggs sometimes started hatching during collection. This was reported on the filed forms and such nests were excluded from the picture analysis.

#### Total brood area

Using ImageJ (Schindelin et al. 2012), the total surface of the brood was measured in number of pixels, by following the outline of the brood as closely as possible. Using the surface of one square from the grid pattern printed on the inside of the nest, the brood surface was converted into a unit comparable across nests (Supplementary File F2, Part 1.3 and 1.4).

#### Cannibalism index

Females lay eggs as tightly packed as possible. Thus, clear empty spaces within the brood are unlikely to result from egg laying patterns. For a guarded nest, they more likely result from parental cannibalism by the male, although they could also be the result of predation. We estimated the prevalence of cannibalism in a nest by measuring the combined surface of the ten largest empty areas within the previously defined brood area (Supplementary File F2, Part 1.5).

#### Overgrowth index

Similarly, to estimate growth of parasites detrimental to the brood within the brood area, we measured the combined surface of the ten largest overgrown areas within the brood (Supplementary File F2, Part 1.6).

#### Egg density and reproductive success

For each brood, a square mask was created in ImageJ, containing approximately between 50 and 100 eggs. The area of the mask in pixels was measured, and the number of eggs within the mask counted. The mask was then moved to another location on the brood and the counting repeated, five times (Supplementary File F2, Part 1.7). This allowed us to estimate the total number of eggs (reproductive success) by extrapolating to total the brood surface, as well as to estimate egg density.

#### Developmental stage

All eggs in the same nest usually hatch within 24h, even if they were laid by different females (Mobley et al. 2009, de Jong 2011). Nevertheless, when the developmental stage is close to the hatching date, empty spaces within the brood could be due to early hatching rather than cannibalism. We took note of nests that appeared close to hatching (clear eggs with very visible eyes) in order to exclude these nests from the analysis of cannibalism index. Such nests were kept in the analysis of brood area (Supplementary File F2, Part 1.8).

### 8. Picture analysis of egg size

The contents of 186 nests preserved in 95% ethanol during the collection of 2022 were analysed in February 2025. The nest acetate sheets and eggs were laid flat under a stereomicroscope (Leica MZ 95) and photographed with an integrated Leica MC170 HD camera and the LAS X software. A microscopic slide with engraved scale was used to set the scale of the images. Pictures were analysed in imageJ: two orthogonal diameters were measured on 5 randomly picked eggs for each nest. The detail of the egg picture analysis protocol is given in Supplementary File F2, Part3.

### 9. Statistical analysis

We describe first (Section 9.1) the analysis of the effect of the thermal environment on aspects of the reproductive dynamics other than reproductive success (i.e. egg number): proportion of occupied nests, egg density and egg size in occupied nests. Then we describe the path analysis of reproductive success (Section 9.2), and finally the post-hoc analysis of the effect of male size on reproductive success (Section 9.3).

#### 9.1 Nest occupancy, egg density and egg size

We assessed the effects of thermal environment (DDF index), time period and nest size on (i) the presence of live eggs, (ii) egg density in the nest, and (iii) egg diameter using three different models. The general approach was to fit a linear mixed-model with sampling site as a random effect, and the fixed effects of DDF, time period, nest size and all possible interactions. Interactions that were not statistically significant were excluded, and a type 2 ANOVA table was reported if no interactions were present in the final model, type 3 otherwise. The presence of live eggs in the nest was analyzed with a generalised linear mixed-model with binomial response (base R, R core team 2023). The other two models assumed a normally distributed response.

#### 9.2 Path analysis

We used a path diagram to describe the hypothesised causal relationships between reproductive success (number of eggs), thermal environment (DDF index), male traits (length and condition), nest quality (size of artificial nest) and nest status (cannibalism and overgrowth in the brood). Our primary interest was to understand how the thermal environment, which varied with latitude, impacted the entire system and ultimately the per-brood individual reproductive success of males in the different populations. Thus, we primarily investigated among-population patterns; however, reproductive success is also likely to depend on local processes occurring within populations, such as competition among males and female mate choice. For this reason we decomposed the male traits and nest status variables into an among-population component (population mean) and within-population component (individual deviation from population mean). When describing model structure, effects referred to as “mean” represent among-population effects, while “relative” effects represent within-population effects. All numerical variables were scaled to a mean of zero and variance of 1, so effect sizes are comparable across models. The structure of the path diagram (Supplementary Figure S2) represents our expectation of how the variables interact in the system: each box is a variable and each arrow represents a possible causal relationship. To quantify and describe these relationships, we use 6 statistical models that are represented graphically in the diagram (Supplementary Figure S2). Each black circle, numbered from 1 to 6, marks the response variable in a statistical model, and each arrow pointing to a circle represents a fixed effect in the model. The structures of the models are described in detail below. Each of the six models was fitted to two separate datasets, corresponding to peak season and late season sampling. The spatial structure of the data was accounted for by including the sampling site as a random effect in each model. Finally, collinearity between fixed effects was assessed by computing variance inflation factors with the *vif* function from the *car* package (Fox and Weisberg 2019), which led to the exclusion of some effects in some models, detailed below (VIF threshold 10, see below).

#### Model 1: Effectsh on individual reproductive success

The response variable, estimated total number of eggs in the brood, was assumed to be normally distributed (count data with very high mean, and after visual check of the distribution). We used the *lme4* package (Bates et al. 2015) in R to fit a linear mixed-effect model, where the sampling site was fitted as a random effect. Fixed effects in the initial model comprised (i) among-population effects: DDF (thermal index), male mean length, male mean condition, mean cannibalism index, mean overgrowth index, and (ii) within population effects: nest size, relative male length, relative male condition, relative cannibalism index and relative overgrowth index.

##### Peak season model

Mean male condition was dropped from the model due to collinearity with DDF, following recommendations from Zuur et al. (2010).

Late season model: Mean overgrowth index was dropped from the model due to collinearity with DDF.

#### Model 2: Effects on cannibalism index

Because the response variable was a true proportion data (surface of the brood cannibalised), we fitted a mixed effect beta regression as implemented in the *glmmTMB* R package (Brooks et al. 2017). There were very few zeroes, consequently we transformed the data as proposed by Smithson and Verkuilen (2006) and proceeded with a beta regression. The transformation is done as follows:

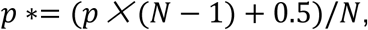

where N is the sample size, p is the proportion data and p* is the transformed data.

Fixed effects in the initial model comprised (i) among-population effects: DDF index, male mean length, male mean condition, and (ii) within-population effects: nest size, relative male length, relative male condition.

Peak season model: Mean male condition was dropped from the model due to collinearity with DDF.

#### Model 3: Effects on overgrowth index

Overgrowth index was also a true proportion data, but in this case with a substantial proportion of zeroes. We resorted to a hurdle model (also sometimes referred to as “zero-inflated beta regression”), where we modelled separately the occurrence of zeroes and the positive values. The first model is a generalized mixed effects model where the binomial response variable is presence/absence of overgrowth. The second is a beta regression as for the case of cannibalism, analysing only the subset of the data with strictly positive values. In both models, fixed effects comprised (i) among-population effects: DDF index, male mean length, male mean condition, and (ii) within-population effects: nest size, relative male length, relative male condition.

Peak season model: Mean male condition was dropped from the model due to collinearity with DDF.

#### Model 4: Effects on nest quality

We provided in all populations standard artificial nests of the same two sizes, either large or small. Thus, we fit a generalized mixed-model with a binomial response variable corresponding to the nest size category. Fixed effects comprised (i) among-population effects: male mean length, male mean condition, and (ii) within-population effects: nest size, relative male length, relative male condition.

#### Model 5: Effect of thermal environment on male length

We fitted a linear mixed effects model with male body length as the response variable and DDF index as a fixed effect.

#### Model 6: Effect of thermal environment on male condition

We fitted a linear mixed effects model with male body length as the response variable and DDF index as a fixed effect.

### 9.3 Post-hoc model

To further investigate the effects of male relative length (deviation from population mean) and nest size on reproductive success, a model was fitted subsequently to the path analysis. In this linear mixed-effects model, the response variable was reproductive success, sampling location was fitted as a random effect, and nest size, time period and male relative length were fixed effects. In addition, the only significant interaction of nest size and male relative length was kept in the final model.

## Results

### 1. Nest occupancy across populations and sampling periods

The probability for an artificial nest to contain a live brood (combined red and orange categories in Figure 3) was higher in peak breeding season than in late season (Peak season: Estimate=0.57, SE=0.043; late season: Estimate= 0.41, SE=0.032, see ANOVA in Table 1). The probability to contain a live brood also increased with DDF (Table 1), from a probability of 0.40 (SE=0.05) at high latitude to a probability of 0.58 (SE=0.06) at low latitude. However, the DDF effect was driven solely by the high occupancy in Kristineberg peak season, and may not in fact be due to the thermal environment. The probability of containing a live brood was not affected by the size of the artificial nest, nor by interaction of nest size and DDF or nest size and time period (Table 1). In the remainder of the result section, we focus on nests containing live broods.

**Figure 3.**
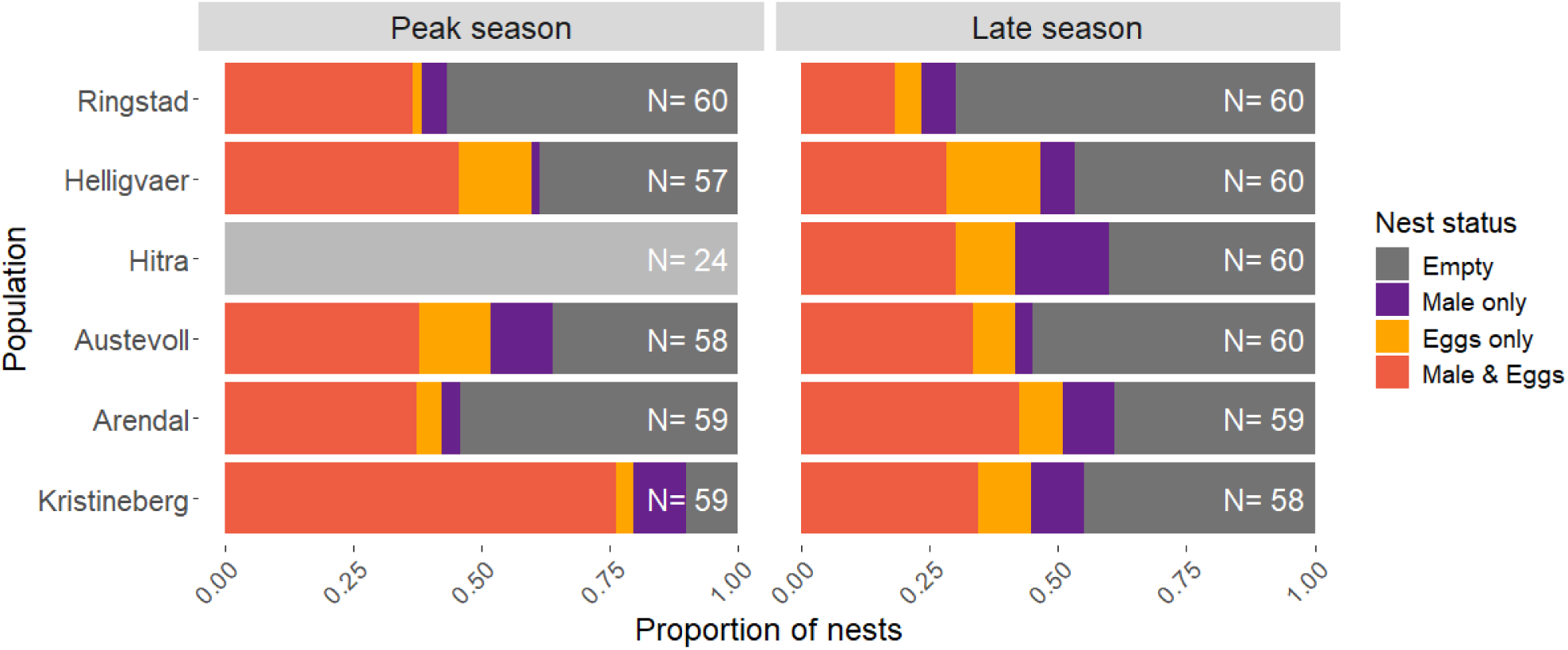
Occupancy status of artificial nests placed in the field for the six P. flavescens populations and two time periods. Each panel corresponds to a time period (Peak season=June 2022, Late season=July 2022). The populations are ordered from north (top) to south (bottom). Each bar gives proportions of nests, with the sample size indicated on the bar. The presence of a guarding male was assessed by a snorkeling observer. The presence of live eggs was assessed on land by extracting the inside of the nest. Nests marked “Eggs only” contained live eggs but no male could be observed entering the nest during the observation period. Peak season Hitra was excluded from the nest occupancy analysis due to a storm that interfered with the collection protocol.

**Table 1.**
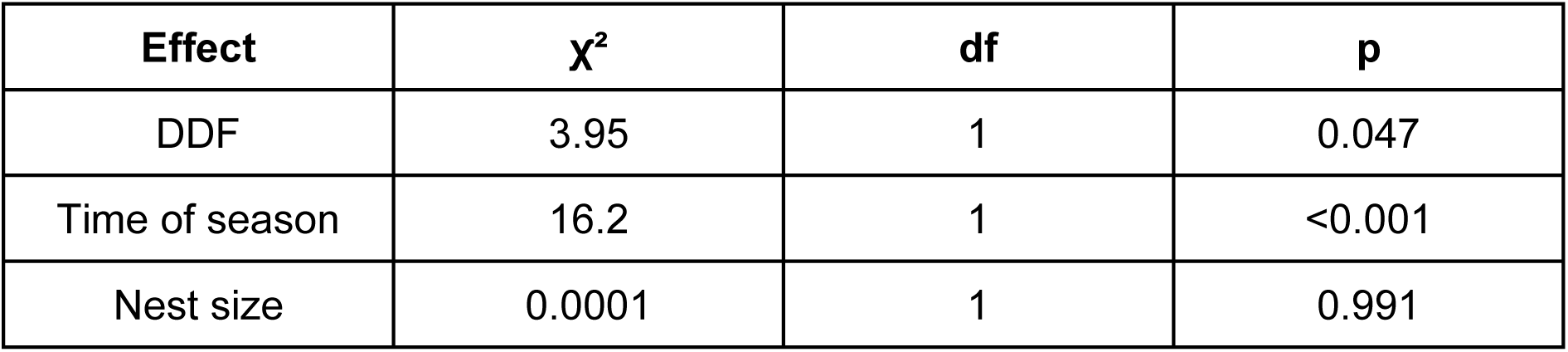
Analysis of variance (type II) for the binomial generalised linear mixed-model with presence of live eggs as a response variable.

### 2. Effect of the thermal environment on males and their broods and drivers of individual reproductive success

We use a path diagram to visualise the relationships between the thermal environment (DDF index), traits of nest-guarding males (body length and body condition), nest status (cannibalism and overgrowth indices), nest quality (artificial nest size), and their combined effects on individual reproductive success. We present two diagrams based on the same structure, one for peak breeding season data (June 2022, Figure 4a) and one for late breeding season data (July 2022, Figure 4b). Each diagram is informed by 6 statistical models, and the ANOVA tables for these models are provided in Supplementary tables 1 to 12. Importantly, our analysis allows for both among-population effects and within-population effects (Methods 8.2). The thermal environment (DDF) has only an among-population component, since there is only one thermal environment value for all individuals in the same populations. Male and nest traits (male length and condition, overgrowth and cannibalism indices) have both an among-population component (mean value of the population) and a within-population component (individual deviation from population mean). Nest quality has only a within-population component since the same exact nests were used in all locations (no variation in population mean). Among-population effects are represented on the diagrams by black arrows and within-population effects by yellow arrows (Figure 4).

**Figure 4.**
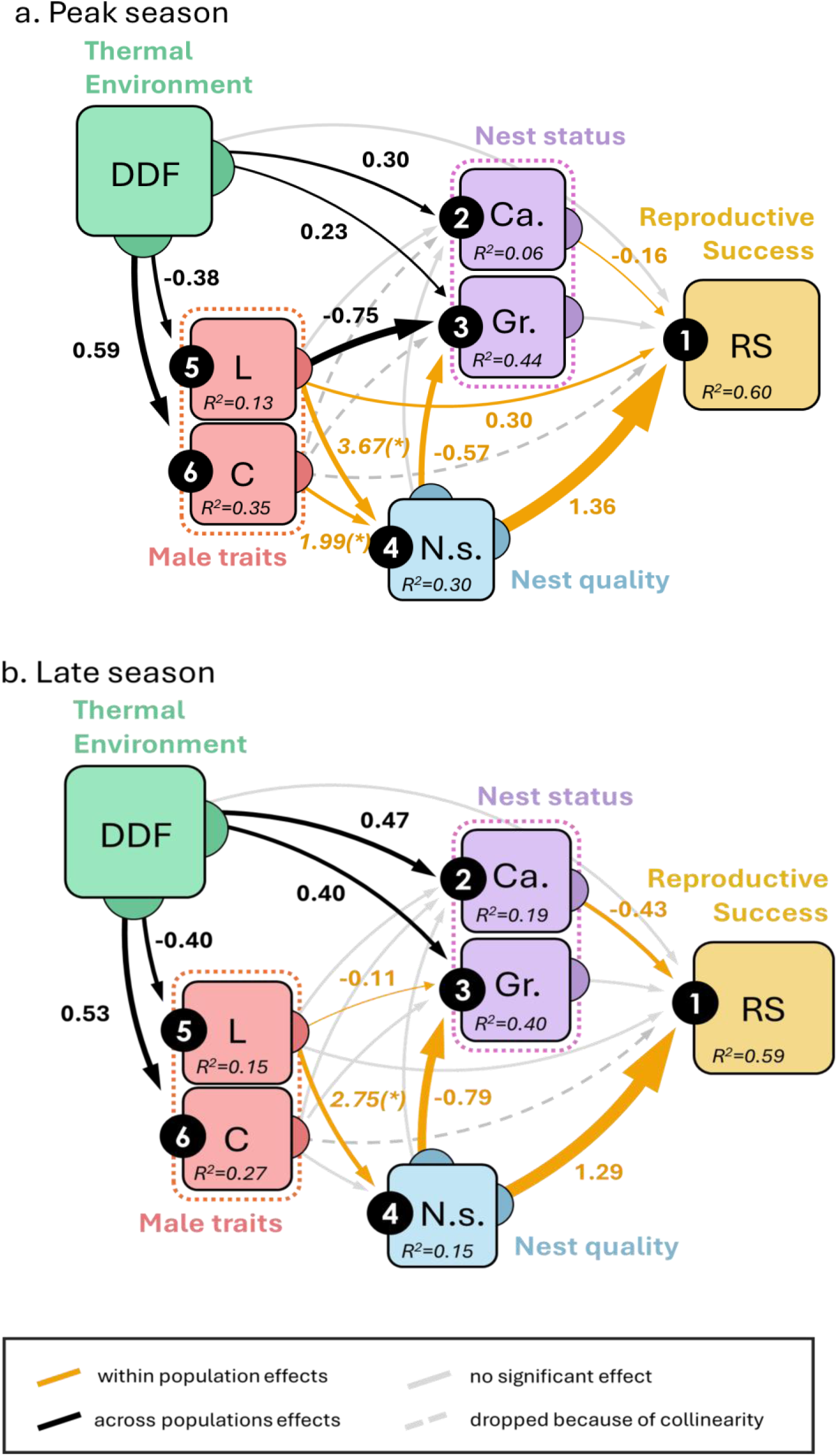
Path diagram summarizing the relationship between thermal environment, male traits, nest status variables, nest quality and reproductive success for six populations of Pomatoschistus flavescens in peak breeding season (a) and late breeding season (b). Each box in the diagram is a variable and each black circle is the response variable in a statistical model. Each arrow pointing to a black circle corresponds to an effect in the model. Black arrows (among-population effects) and yellow arrows (within-population effects) represent statistically significant effects (threshold p=0.05). Thin grey arrows represent effects tested but not significant. The thickness of the arrows is proportional to effect size, reported next to the arrow. All numerical variables are scaled so that effect sizes represent a change in units of standard deviation of the response variable, caused by a 1 standard deviation change in the explanatory variable. For nest size as an explanatory variable: the effect size is the change in units of standard deviation of the response variable, caused by changing from small to large nest. (*) For nest size as a response variable in model 4: effect size is the change in odds ratio to obtain a large nest, for a change of 1 standard deviation in the explanatory variable. The ANOVA tables for the models are given in Supplementary tables 1 to 12.

The thermal environment (DDF index) affected male traits and nest status variables consistently across the two sampling periods. However, male reproductive success was not affected by the thermal environment, but instead responded exclusively to within-population effects with some differences between peak season and late season (Figure 4a and 4b). We explore these patterns in detail in the next two sections.

#### 2.1 Effects of the thermal environment on male traits and nest status

The thermal environment had consistent effects across time periods on male traits (length and body condition) and nest status (cannibalism and overgrowth indices, Figure 4a and 4b). Body size of breeding males tended to increase and body condition to decrease as the thermal environment got more challenging (lower DDF, higher latitude, Figure 5). Overgrowth in the nest and brood cannibalism both tended to decrease in more challenging thermal environments (lower DDF, higher latitude, Figure 6).

**Figure 5.**
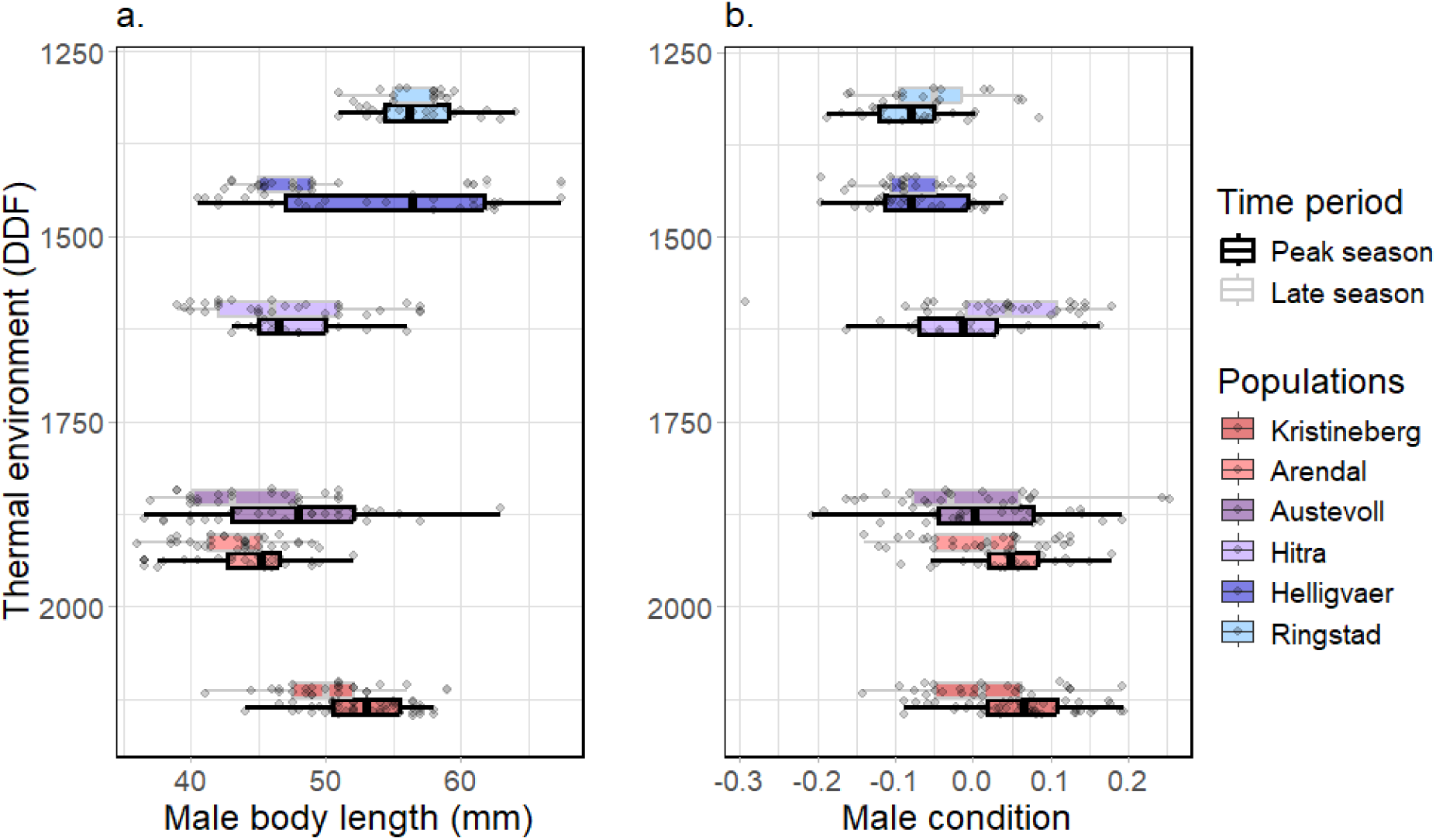
Body length (a) and body condition (b) of nest-guarding males across six populations of P. flavescens experiencing different thermal environments. The y-axis of both panels show the thermal environment index (DDF) in unit of degree days, with the northern populations at the top (lowest DDF). Points are jittered along the y axis for better visualisation, but each population has only one fixed value of DDF (see Figure 2).

**Figure 6.**
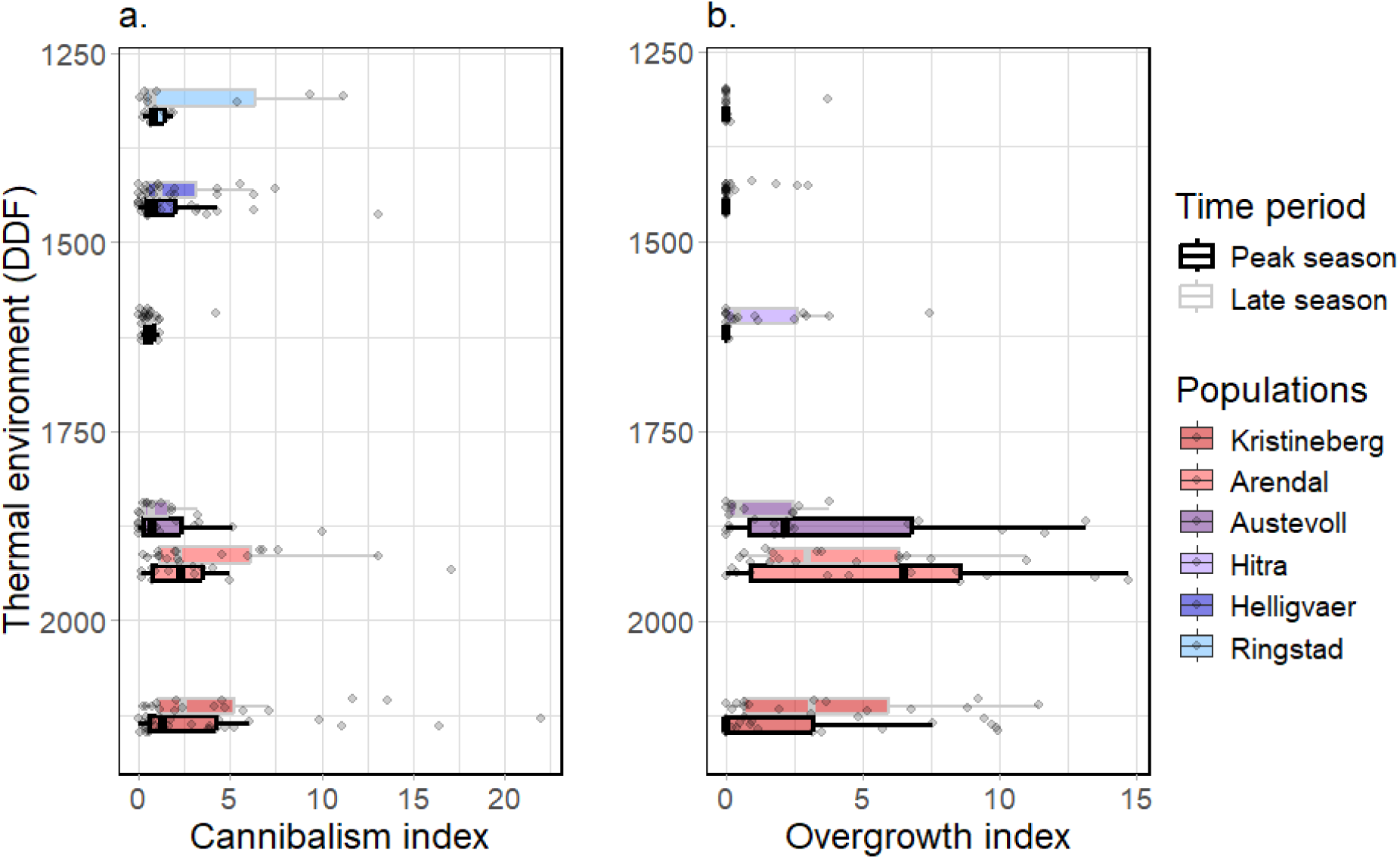
Cannibalism index (a) and Overgrowth index (b) of occupied nests across six populations of P. flavescens experiencing different thermal environments. The y-axis of both panels show the thermal environment index (DDF) in units of degree days, with the northern populations at the top (lowest DDF). Points are jittered along the y axis for better visualisation, but each population has only one fixed value of DDF (see Figure 2).

The brood cannibalism index was not affected by either nest quality or male traits (Figure 4a and 4b). In contrast, the overgrowth index was affected by nest quality, with larger nests having less overgrowth, and by male length with larger males having less overgrowth in their nests. More specifically, male mean length had an effect in peak season (Figure 4a), and male relative length in late season (Figure 4b). The effects of nest size and male size on the overgrowth index are displayed in Supplementary figure S3.

#### 2.2 Within-population effects on reproductive success

Across both time periods, the largest effect on reproductive success was caused by nest quality (size of artificial nest), which was itself driven by the body length of the guarding male relative to the population mean (Figure 4a and b). The effect of nest size on reproductive success was substantial with roughly a doubling of clutch size from small to large nests (Peak season averages: small nests= 5714, SE=746; large nests= 11670, SE= 804; Late season averages: small nests= 4879, SE= 727; large nests= 10043, SE=699). A relative increase of 1 standard deviation in length (equivalent to 5 mm in peak season, 4.7 mm in late season) increased the odds ratio to occupy a large nest by 3.7 times and 2.8 times in peak season and late season respectively. Reproductive success was affected negatively by the amount of cannibalism within populations but the global impact was small (see Supplementary Figure S4 for the relationship between relative cannibalism index and reproductive success).

During the peak of the breeding season but not in late season, male relative length contributed directly to reproductive success, in addition to the indirect contribution through acquisition of larger nests (Figure 4a). This direct effect had a smaller effect size: an increase of 1 standard deviation in relative length translated into 1297 additional eggs. Still in peak season, the acquisition of larger nests was driven by male condition relative to the population mean, in addition to male relative length (Figure 4a). The within-population effects of cannibalism, nest size and male traits combined explained about 60% of the variation in reproductive success at both time periods (see Supplementary figure S5 for the reproductive success per population and time period).

We explored further the relationship between reproductive success and relative male length for each nest size and time period (Figure 7). A post-hoc model fitting the effect of male relative length, nest size, time period and their interactions on reproductive success was used to better describe this complex system (ANOVA in Table 2). The relationship between male relative length and reproductive success estimated from this model indicated a significantly positive slope in large nests (Peak season: Slope=295 Eggs/mm, SE=67.3, p<0.0001; Late season: Slope= 236 Eggs/mm, SE=89.2, p=0.017), but no significant relationship between male relative length and reproductive success in small nests.

**Figure 7.**
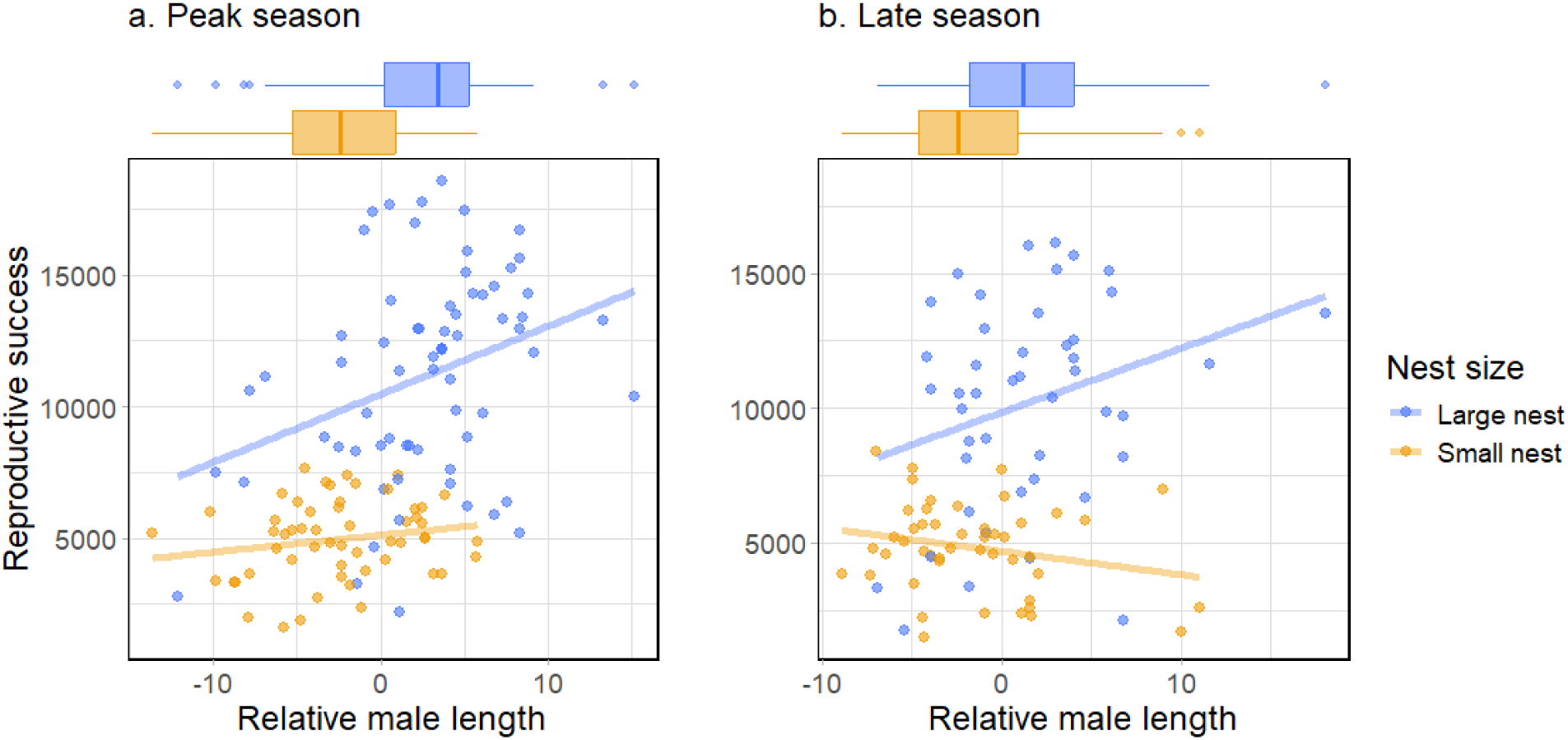
Reproductive success in P. flavescens as a function of relative male length, during (a) peak breeding season and (b) late breeding season . Each data point corresponds to a nest and its guarding male, points are colored by nest size and trend lines corresponding to a linear regression are shown for each time period and nest size combinations. Male length is made relative by subtracting the population mean. The marginal boxplots above each panel indicate the distribution of relative length of guarding males in large nests (blue) and small nests (yellow) for each time period.

**Table 2.**
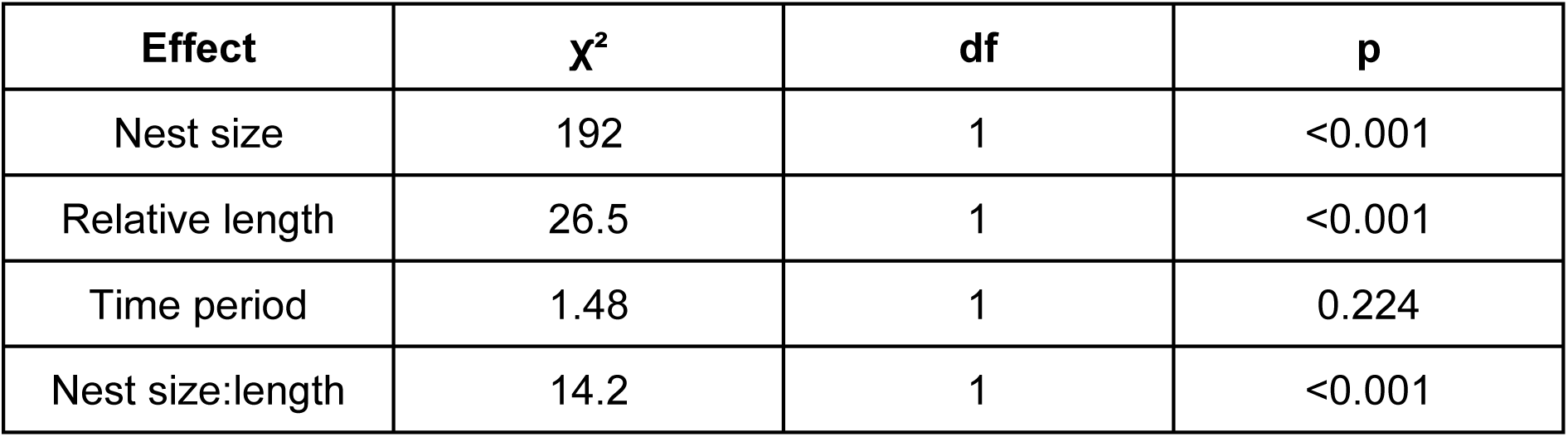
Analysis of variance (type III) for the linear mixed-model with reproductive success as a response variable.

| Effect | $\chi^2$ | df | p |
| --- | --- | --- | --- |
| Nest size | 192 | 1 | <0.001 |
| Relative length | 26.5 | 1 | <0.001 |
| Time period | 1.48 | 1 | 0.224 |
| Nest size:length | 14.2 | 1 | <0.001 |

### 3. Effect of the thermal environment on egg density and egg size

We were primarily interested in individual reproductive success, as measured by the number of eggs in a nest. However, important information on the reproductive strategy can be gained from assessing egg sizes and density. Egg density was measured within the brood area, and thus reflects how tightly packed the eggs were within the brood, not how full the nest was.

As the thermal environment got more challenging (lower DDF, higher latitude), egg density in the nests decreased by about 18% (Figure 8a, Table 3), while egg diameter increased by about 14% from South to North (Figure 8b, Table 4). However, egg density variation was not fully constrained by egg size, since small nests had overall higher egg density than large nests, irrespective of latitude (Table 3, estimated means: Small nets=188 eggs/cm^2^, SE=3.02; Large nests=166 eggs/cm^2^, SE=2.97). Finally, egg diameter was generally higher during peak breeding season as compared to late season (Figure 8b, Table 4, estimated means: Peak season=0.70 mm, SE=0.0054; Late season=0.67 mm, SE=0.0053).

**Figure 8.**
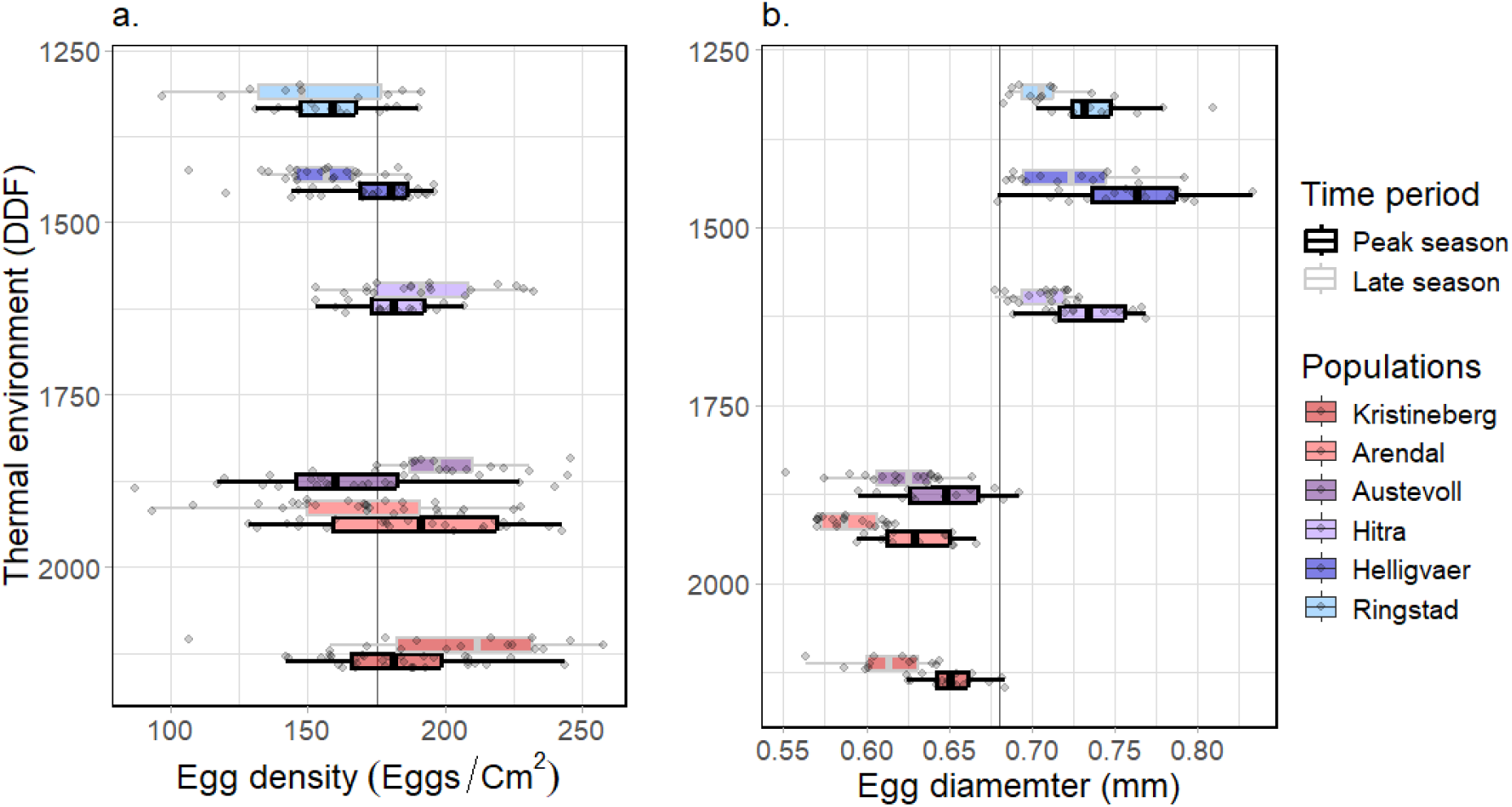
Egg density (a) and egg diameter (b) in artificial nests across six populations of P. flavescens experiencing different thermal environments. The y-axis of both panels show the thermal environment index (DDF) in units of degree days, with the northern populations at the top (lowest DDF). Points are jittered along the y axis for better visualisation, but each population has only one fixed value of DDF (see Figure 2). Vertical grey lines are drawn to help comparison between populations.

**Table 3.**
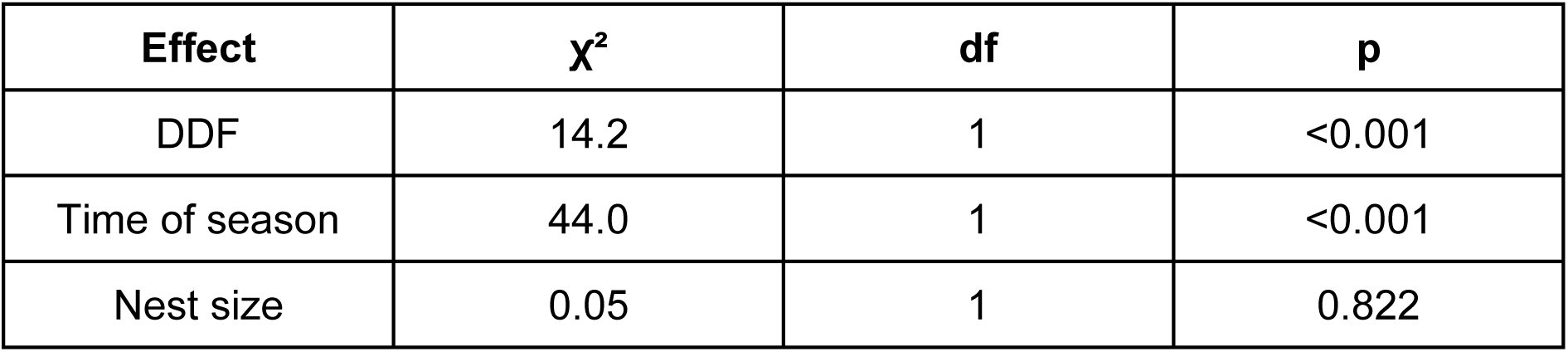
Analysis of variance (type II) for linear mixed-model with egg density as a response variable.

| Effect | $\chi^2$ | df | p |
| --- | --- | --- | --- |
| DDF | 14.2 | 1 | <0.001 |
| Time of season | 44.0 | 1 | <0.001 |
| Nest size | 0.05 | 1 | 0.822 |

**Table 4.** Analysis of variance (type II) for linear mixed-model with egg diameter as a response variable.

| Effect | $\chi^2$ | df | p |
| --- | --- | --- | --- |
| DDF | 69.4 | 1 | <0.001 |
| Time of season | 1.62 | 1 | 0.203 |
| Nest size | 64.17 | 1 | <0.001 |

## Discussion

We studied nest-brooding males of *Pomatoschistus flavescens* in six natural populations across a steep climatic gradient, to gain insights on the ecological sensitivity of reproductive dynamics in coastal ecosystems of Scandinavia. The core result was that climate, represented by the thermal environment in the breeding habitat, did not affect the individual reproductive success of breeding males. This finding was intriguing considering the impact that climate has on other aspects of the reproduction of *P. flavescens*. Male phenotypes and brood cannibalism, important determinants of reproductive success, were affected by climate. Timing of reproduction and population density (Martinossi-Allibert et al. 2025a), as well as operational sex ratio (Martinossi-Allibert et al. 2025b), are also affected by climate. Moreover, brood development speed is temperature-dependent (Skolbekken and Utne-Palm 2001). Thus, with the use of standardized nests, we expected that climatic effects on male size, development speed, cannibalism and population sex ratio would carry over to affect the number of eggs guarded by each male. Yet, and despite a steep climatic gradient, we detected no such effect, which suggests the mediation of a buffering mechanism between ecological conditions and reproductive success. In turn, we found that reproductive success was largely determined by the deviation of individual male phenotypes from the local average, highlighting the role of mating competition. We raise the possibility that local competition, a form of frequency-dependent or soft selection, may buffer reproductive success against variation in climatic conditions. If general, such mechanisms could have important implications for the resilience of natural populations to environmental change.

Frequency dependence plays a prominent role in evolutionary processes (Wright 1931, Svensson 2018, Gómez-Llano et al. 2024), but it is usually not considered in studies on adaptation to environmental change and evolutionary rescue (Svensson and Connallon 2019, Araya-Ajoy et al. 2025a). This omission likely limits the validity of predictive population dynamics in changing environments (Engen et al. 2020, Araya-Ajoy et al. 2025b), but there is in fact a lack of empirical data on frequency dependence interacting with ecological conditions. Across the latitudinal gradient of our study, reproductive success responded to male traits (size, condition) or nest status (cannibalism) in a frequency dependent manner, because it was solely affected by individual deviations from the local mean of these variables, and this in spite of climate-associated variations in the population means of the traits. We consider potential implications of reproductive success being mainly driven by frequency-dependent processes for population demography under environmental change, in *P. flavescens* and more generally.

To begin with, frequency dependence implies a demographic cost, both because of competition and maladaptation. The energy expended by males *P. flavescens* to compete for high quality nests or court females is energy that cannot be allocated to parental care, meaning reduced reproductive success. Furthermore, maladaptation adds to the demographic cost if the traits favored by social competition, colorful ornaments, mating calls and displays, do not contribute to ecological adaptation (Chenoweth et al. 2015, Berger et al. 2016, Martinossi-Allibert et al. 2019b, Svensson an Connallon 2019, McGlothlin and Fisher 2021). Yet, these negative impacts of frequency dependence on demography should be nuanced. First, social selection can in fact contribute to ecological adaptation, for example if the size and ornamentation of males *P. flavescens* are honest signals of thermal adaptation or parasite resistance (e.g. Hamilton and Zuk 1982). Such positive effects may be prominent in stressful environments, where ecological and social selections are more likely to align (Connallon and Hall 2016), and may thus be important for evolutionary rescue. Second, the more a trait is determined by social competition the less it is determined by ecological factors, leading to a buffering of the trait against rapid ecological change. This could be what we observe with the reproductive success of *P. flavescens* across Scandinavian populations, which responds to the local distribution of male phenotypes but not to environmentally driven trait variation. Such a buffering of fitness components from ecological conditions could be rather general, since competitive frequency-dependence is expected in every sexually reproducing population where resources required for reproduction are limited (Clutton-Brock 2009, Janicke 2024). In turn, the buffering of reproductive success could result in the maintenance of demographic output under changing environmental conditions. However, we do not expect this mechanism to provide more than a very limited and short term positive effect on population growth under adverse ecological conditions. Indeed, the buffering effect would be limited to fitness components that are highly frequency-dependent, providing limited protection against population collapse. For example, in *P. flavescens* the fitness proxy is reproductive success measured as a number of eggs in the nest, a metric that is not easily extrapolated into population growth due to the high mortality of pelagic larval stages, and important density regulation up to the next recruitment. While individual reproductive success may be buffered against variation in climate, population growth may not. Finally, traits that are highly buffered against ecological variables are unlikely to contribute to ecological adaptation, reducing the prospect for evolutionary rescue in the long-term (Martinossi-Allibert et al. 2019a).

We are convinced that the absence of a climatic effect on reproductive success represents an important biological process at play in these populations. This result is robust because (i) we detected differences among populations in individual male reproductive success, albeit not related to climate, (ii) we detected effects of climate on other variables in the system, and (iii) we explained a large part of the variation in individual reproductive success in the data. Yet our experimental design comes with limitations that we want to acknowledge. For example, while it is an informative approach to use an existing climatic gradient to simulate effects of ecological change (Nagelkerken et al. 2023), we are studying presumably well-adapted populations. These populations may thus exhibit different reproductive strategies allowing them to cope with latitudinal differences in climate (see Martinossi-Allibert et al. 2025a), which could contribute to the absence of a climatic effect on reproductive success. Moreover, although the thermal environment is an obvious environmental constraint for ectotherms, there are likely other ecological factors at play when it comes to limiting reproductive success, such as the local availability of natural nests, or the density and quality of preys. In that view, the population differences in average reproductive success that we observed could have been caused by a combination of environmental factors not considered in our path analysis. The use of artificial nests could also be seen as a limitation of our study design, by under-representing the natural variation in nest sizes and quality; however, in our view artificial nests present several advantages. Beyond the standardizing of nests across populations, sampling only from a limited range of nest types means that the variation in reproductive success we analyze is likely an underestimation of what could be observed in natural nests. Our results are thus conservative in that respect.

Beyond the effect of climate, studying six populations across a large part of the species range allowed us to deepen our understanding of the reproductive dynamics of *P. flavescens*, a model system for the study of sexual selection (Forsgren et al. 2004, Myhre et al. 2012, Utne-Palm et al. 2015, Amundsen 2018, Martinossi-Allibert et al. 2025b). Earlier research restricted to the population of Kristinberg, Swedish west coast, had shown that both female choice (Borg et al. 2006) and male competitive exclusion from nests (de Jong 2011, Wacker et al. 2014) were limited to the early breeding season. While our result agrees with female choice acting solely early on (direct effect of male traits on reproductive success), the path analysis revealed that larger males still obtained larger broods by holding larger nests late in the breeding season, suggesting that competitive exclusion may be maintained after all. An alternative explanation for this pattern could be that small nests are simply preferred by smaller males, enacting a different reproductive strategy. Smaller nests represent a lower parental expenditure due to the smaller brood size (Skolbekken and Utne-Palm 2001) and offer better protection against predators (Lissåker and Kvarnemo 2006), at the cost of restricted oxygen flow and higher brood infection risk, especially in warm waters. Female choice may also act differently in small nests (e.g., based on acoustic signals, Albouy et al. 2023), since male size was associated with reproductive success only within large nests. Here, we described patterns of sexual selection and reproductive strategies across all populations combined. To go further, larger sample sizes would be required to test for interactions between different variables in the path diagram. For example, strategies related to nest size are likely to interact with latitude, due to the temperature dependency of marine growth and oxygen consumption; the strength of sexual selection on male body size could also explain variation among populations in this trait, in particular the large size of high latitude populations (see also Martinossi-Allibert et al. 2025a).

## Conclusion

Using the natural climate laboratory of the Scandinavian coastline, we studied the ecological sensitivity of the reproductive strategy of *Pomatoschistus flavescens*, a nest-brooding fish ubiquitous in these ecosystems. Our analysis revealed that reproductive success was driven in a large part by local competition, and independent of the thermal environment. This result raises the possibility that reproductive success in *P. flavescens* may be strongly constrained by local competition, so that this proxy of fitness, and perhaps to some extent demography, may be buffered against large climatic changes. In natural populations, intraspecific competition imposes a strong constraint on reproductive output, which may lead to compensatory mechanisms when ecological stressors affect the entire competitive pool. Such effects would not be perceived in experimental setups where individual reproductive output is measured outside of the population context. Our results thus support recent theoretical arguments advocating for the integration of social processes to predict population dynamics under environmental change scenarios. Going further, it will be crucial to investigate the generality of this compensatory phenomenon, and deepen our understanding of its potential consequences. Short-term positive effects for population persistence may occur if relaxed competition compensates for adverse effects of climate on population growth, but long-term effects on population persistence are unclear.

## Acknowledgments

We acknowledge the following funding sources: the Research Council of Norway Marinforsk 294453 to the DYNAMAR project, and the Sparebank MidtNorge gavefond. We are also thankful to the field stations of Kristineberg marine research station, and stations of the Havforskningsinstituttet of Norway at Flødevigen and Austevoll. Finally, we extend our gratitude to the numerous field assistants that made this ambitious sampling design a reality: Claudia Aparicio Estallela, Mathias Nyheim, Vera Løvold, Ida Lie-Nilsen, Dagmar M. Coelle, Paula Schmidtz, Henri Martinossi, Karoline Brudevoll Rognlien, Pauline Mesterson Pitter, Erik Amundsen, Amy Li, and Audun Aas Roseth.

