## Supplementary figures for "Local mating competition, but not climate, drives male reproductive success across a latitudinal gradient in a nest-brooding marine fish"

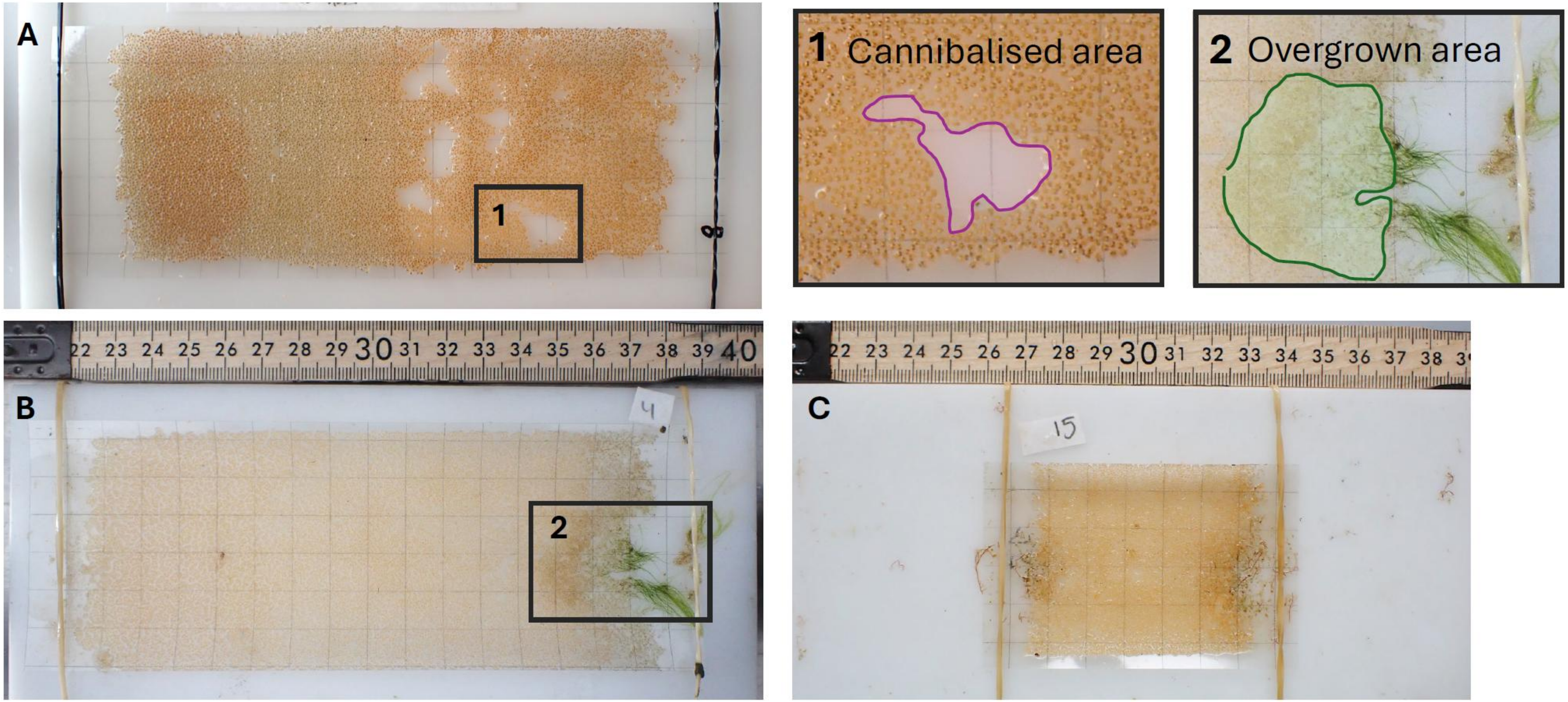


**Supplementary figure S1. Examples of nest pictures.**
Example of nest pictures from the dataset. A,B: Acetate sheets from large nests. C: acetate sheet from a small nest. Panel 1 and 2 show enlargement of areas that would be considered as (1) cannibalized and (2) overgrown. For further details on the picture analysis protocol, see Supplementary file F2.

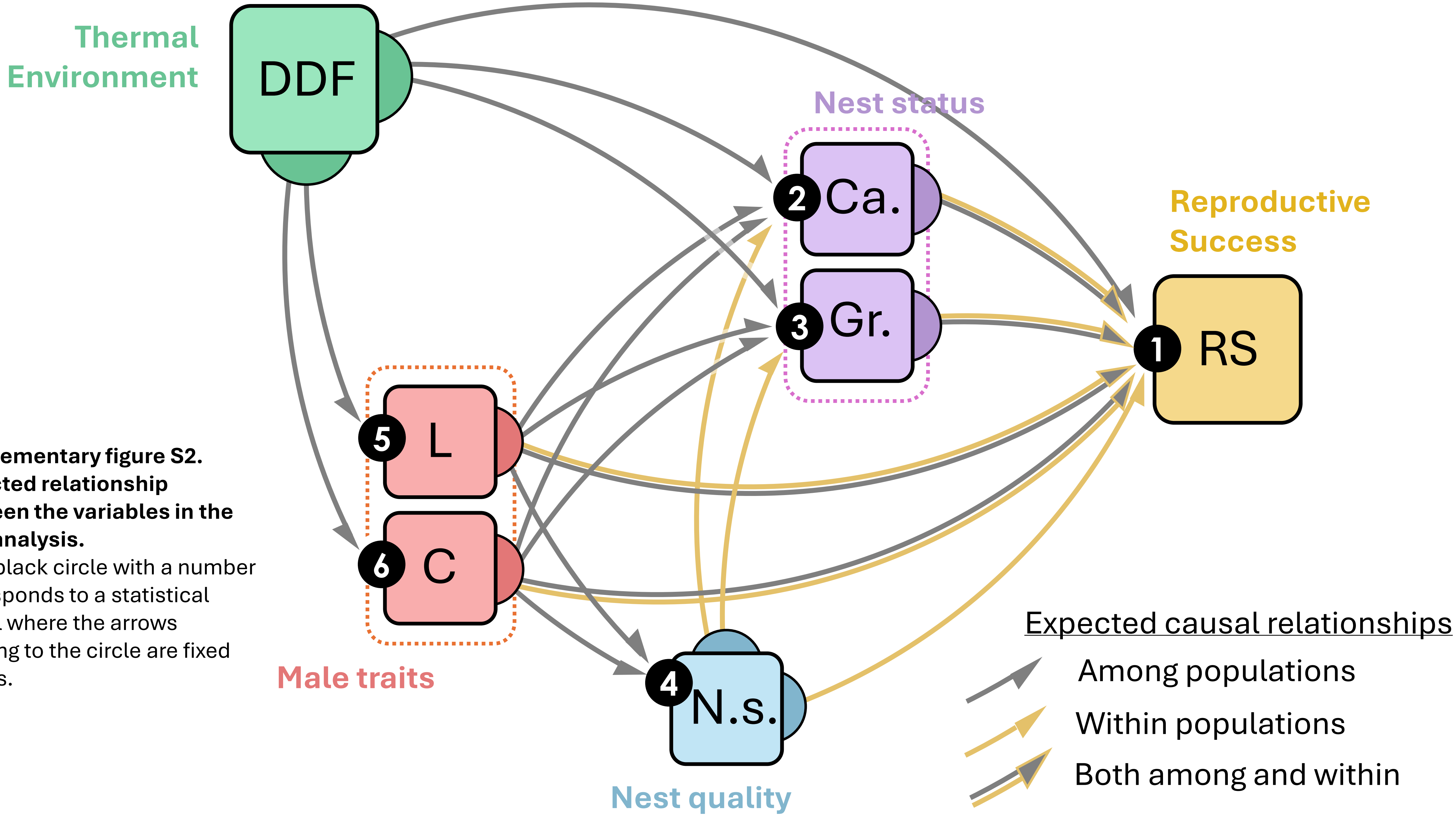


**Supplementary figure S2. Expected relationship between the variables in the path analysis.**
Each black circle with a number corresponds to a statistical model where the arrows pointing to the circle are fixed effects.

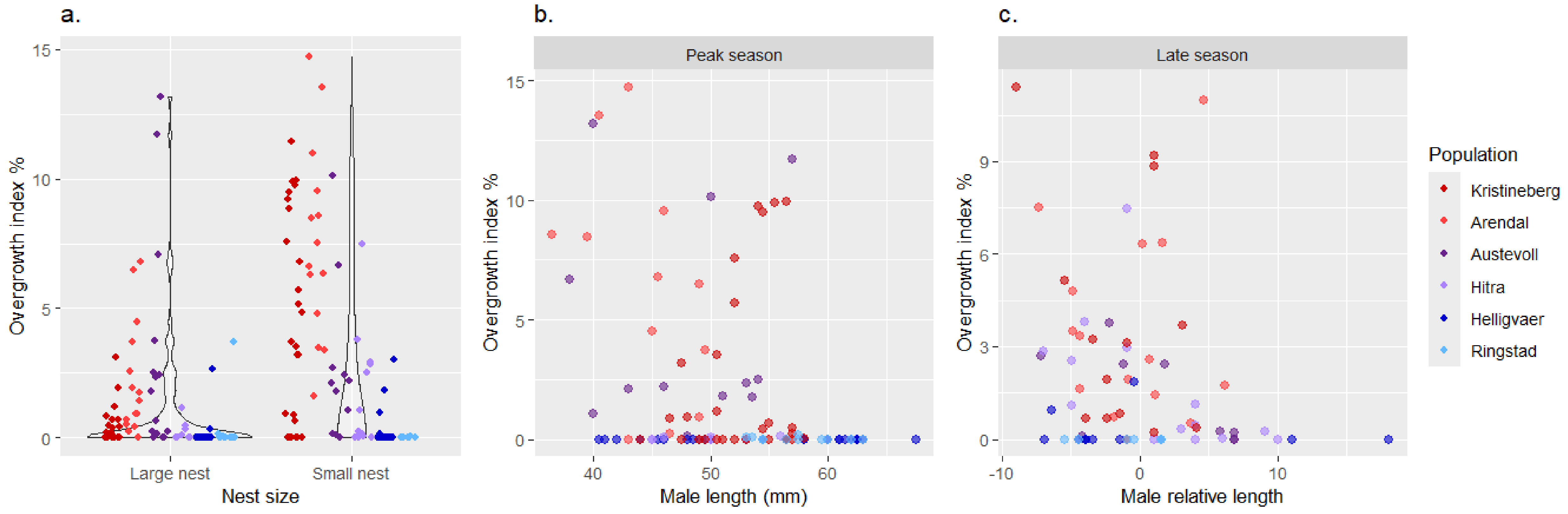


**Supplementary figure S3. Effects of nest size and male length on the brood overgrowth index for the six *P. flavescens* populations.**

The y-axis of all panels shows the overgrowth index, in percent surface of the brood overgrown. In (a), points are jittered along the y axis for better visualization. The color of dots indicates the population from Southernmost (Kristineberg) to Northernmost (Rignstad).

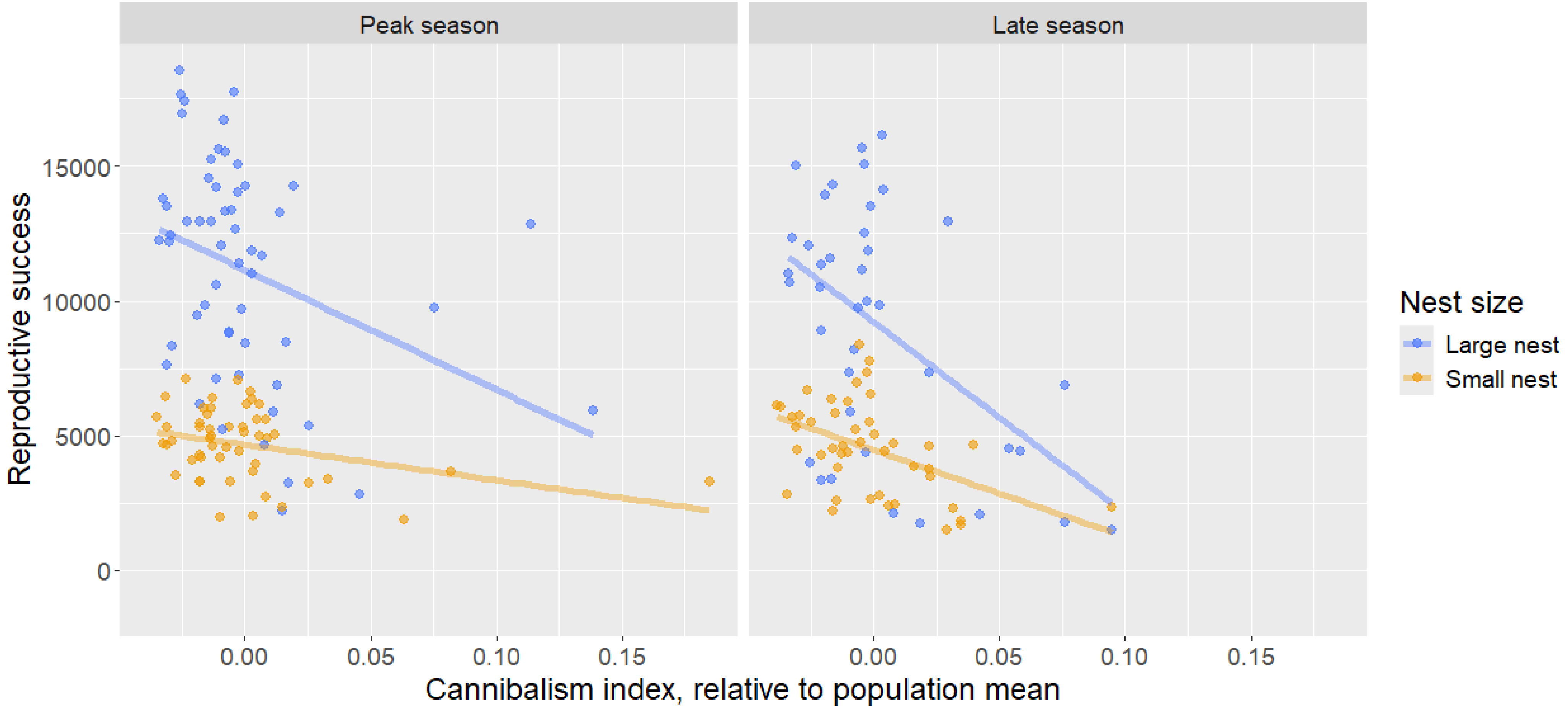


**Supplementary figure S4. Relationship between relative cannibalism index and reproductive success**
Relative cannibalism index is calculated as a difference to the mean cannibalism index of each population.

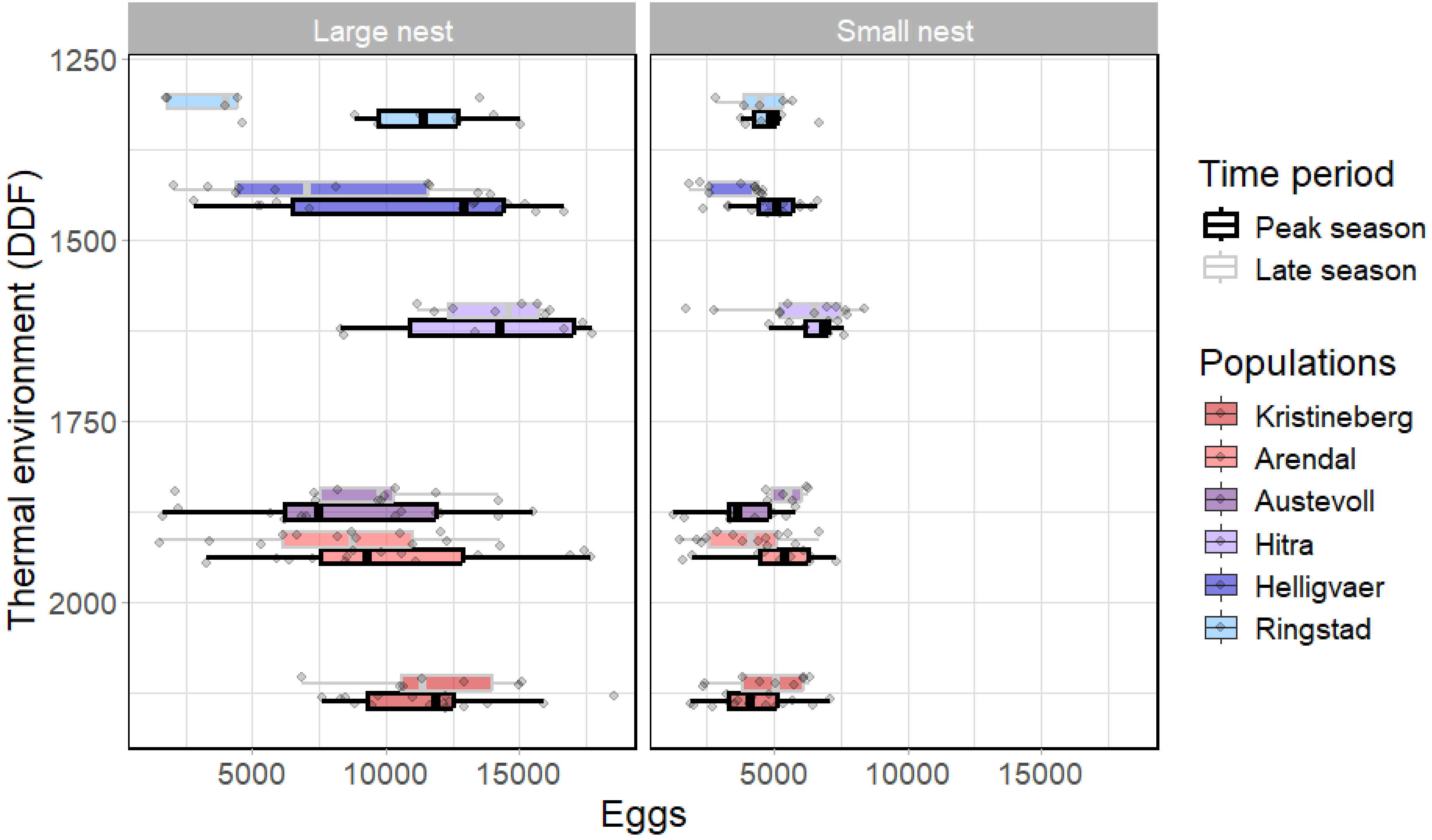


**Supplementary figure S5. Reproductive success for each of the six *P. flavescens* populations, per nest size and time period.** The y-axis of both panels show the thermal environment index (DDF) in unit of degree days, with the northern populations at the top (lowest DDF). Points are jittered along the y axis for better visualisation, but each population has only one fixed value of DDF (see Figure 2).
