## Supplementary file F1 for "Local mating competition, but not climate, drives male reproductive success across a latitudinal gradient in a nest-brooding marine fish"

### Supplemtary file F1 field locations

a. Location of study populations

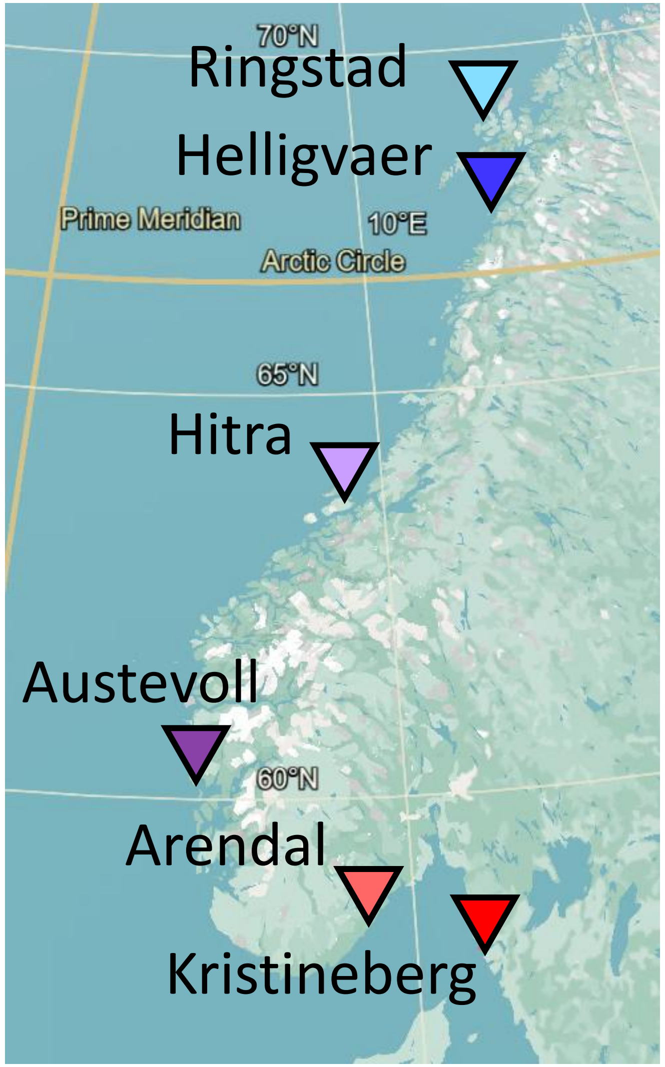


b. Peak season sampling times

| | June | | | | | | | | | | | | | | | | | | | | | | | | | | |
|---|---|---|---|---|---|---|---|---|---|---|---|---|---|---|---|---|---|---|---|---|---|---|---|---|---|---|---|
| | 4 | 5 | 6 | 7 | 8 | 9 | 10 | 11 | 12 | 13 | 14 | 15 | 16 | 17 | 18 | 19 | 20 | 21 | 22 | 23 | 24 | 25 | 26 | 27 | 28 | 29 | 30 |
| Ringstad | | | | | | | | | | | | | | | | | | | DH | | F | | | | | | H |
| Helligvær | | | | | SK | G | | | | LS | | | | | | | | | | | | | | | | | |
| Hitra | | | | | | S8 | F | | T10 | | | | | | | | | | | | | | | | | | |
| Austevoll | | | | | | | | | | | | | | | T | K | L | | | | | | | | | | |
| Arendal | | | | | | | | | | | | | SH | | L | SJ | | | | | | | | | | | |
| Kristineberg | T | | P | B | | | | | | | | | | | | | | | | | | | | | | | |

c. Late season sampling times

| | July | | | | | | | | | | | | | | | | | | | | | | | | | | | |
|---|---|---|---|---|---|---|---|---|---|---|---|---|---|---|---|---|---|---|---|---|---|---|---|---|---|---|---|---|
| | 1 | 2 | 3 | 4 | 5 | 6 | 7 | 8 | 9 | 10 | 11 | 12 | 13 | 14 | 15 | 16 | 17 | 18 | 19 | 20 | 21 | 22 | 23 | 24 | 25 | 26 | 27 | 28 |
| Ringstad | | | | | | | | | | | | | | | | DH | | F | | H | | | | | | | | |
| Helligvær | | | | | | | | | LS | G | SK | | | | | | | | | | | | | | | | | |
| Hitra | | | | | | T10 | | | | F | S8 | | | | | | | | | | | | | | | | | |
| Austevoll | | | | | | | | | | | | | | | | | | | T | L | K | | | | | | | |
| Arendal | | | | | | | | | | | | | | | | | | | SH | SJ | L | | | | | | | |
| Kristineberg | | | | | | | | | | B | T | P | | | | | | | | | | | | | | | | |

The general location of the six study populations is given in (a), and the sampling date (year 2022) for each study site (3 per population) in (b) for peak season and (c) for late season. The sampling sites are color coded by population and the letter corresponds to the name of the samplinmg site, given on the next slides for each population.

#### Transect coordinates for the 3 sub-locations of the Kristineberg population.

Red lines indicate artificial nests lines. 20 nests are placed along each line.

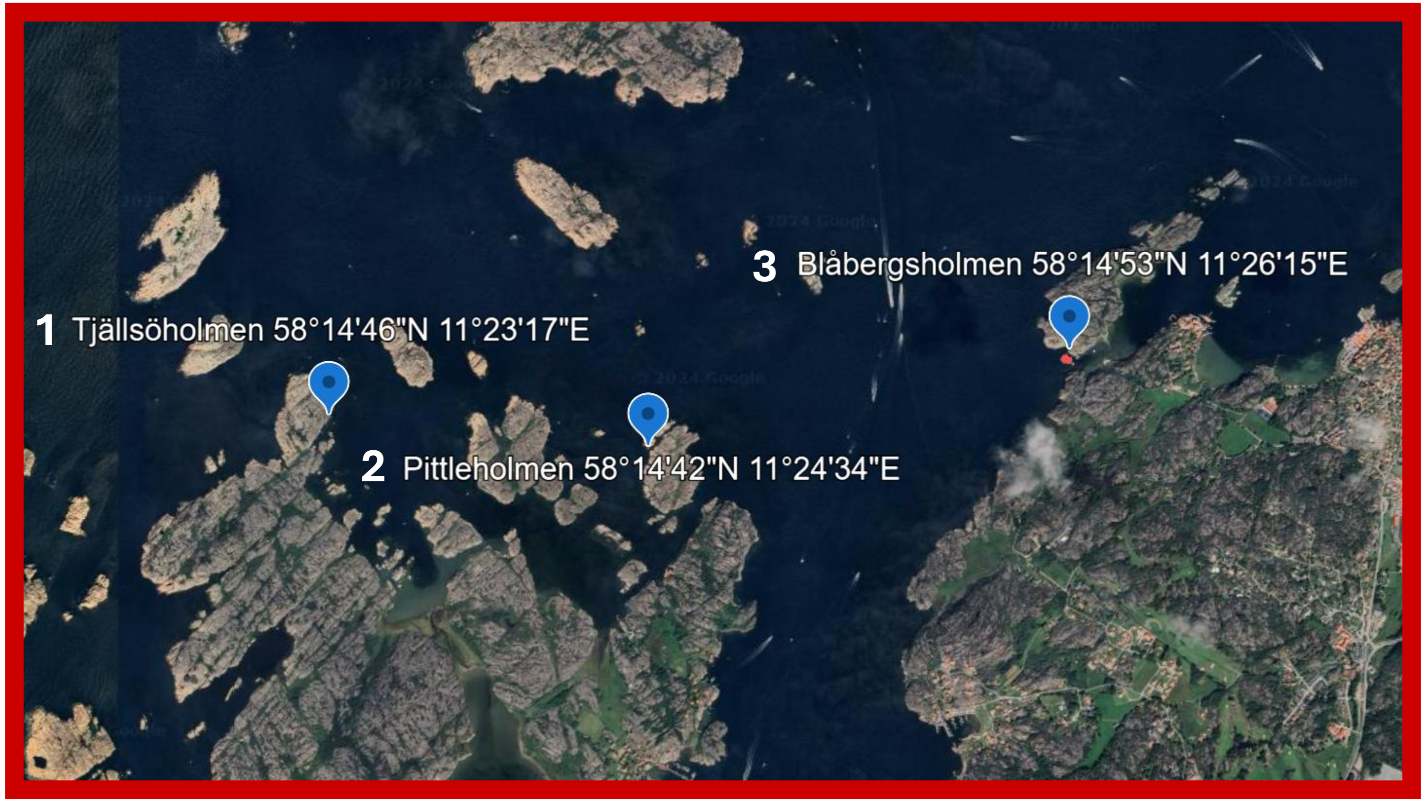


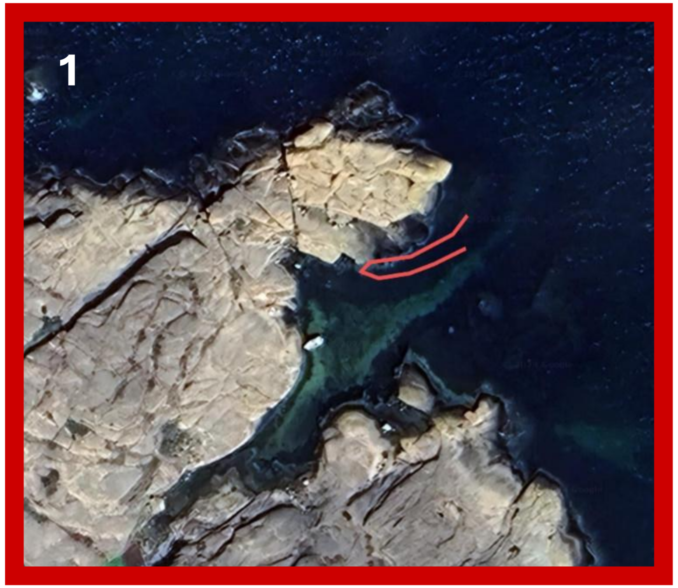


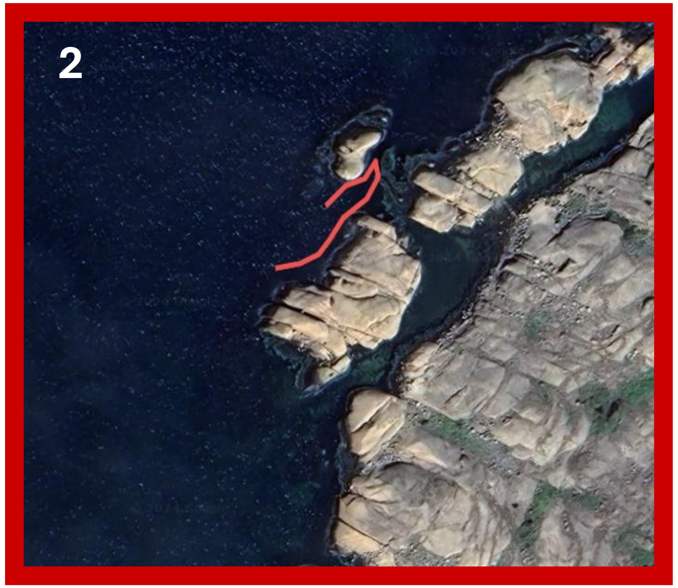


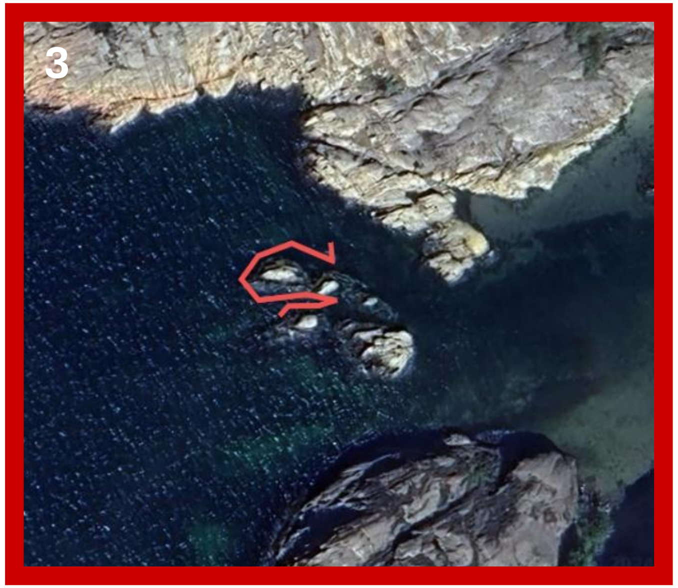

#### Transect coordinates for the 3 sub-locations of the Arendal population.

Red lines indicate artificial nests lines. 20 nests are placed along each line.

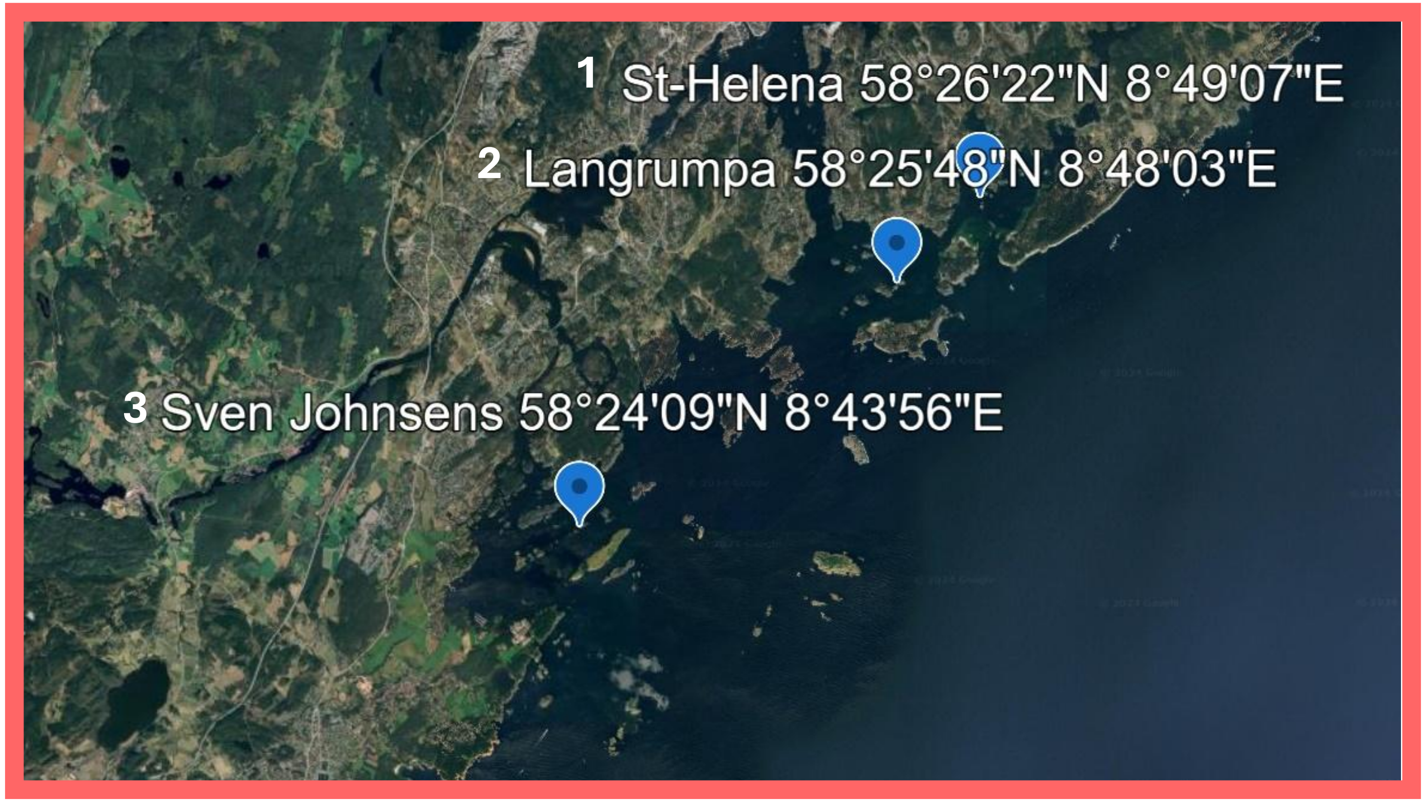


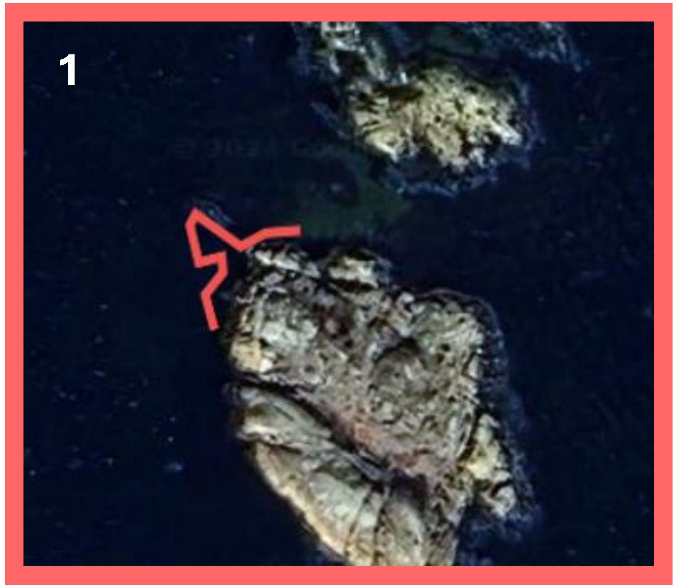


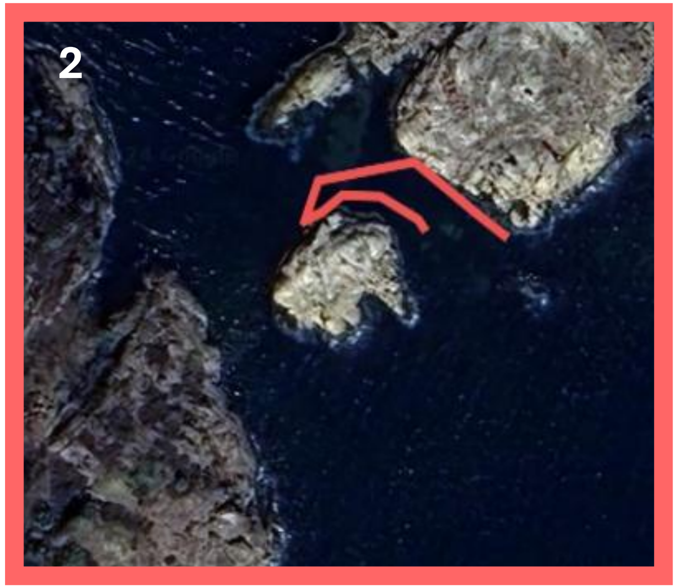


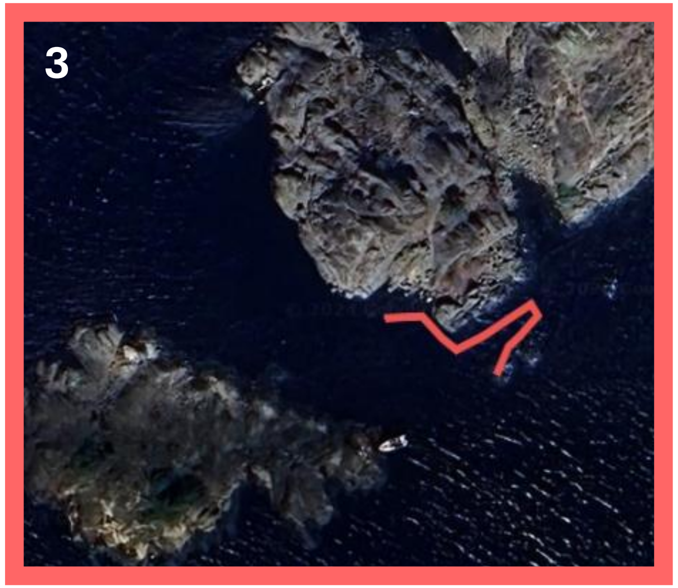

#### Transect coordinates for the 3 sub-locations of the Austevoll population.

Red lines indicate artificial nests lines. 20 nests are placed along each line.

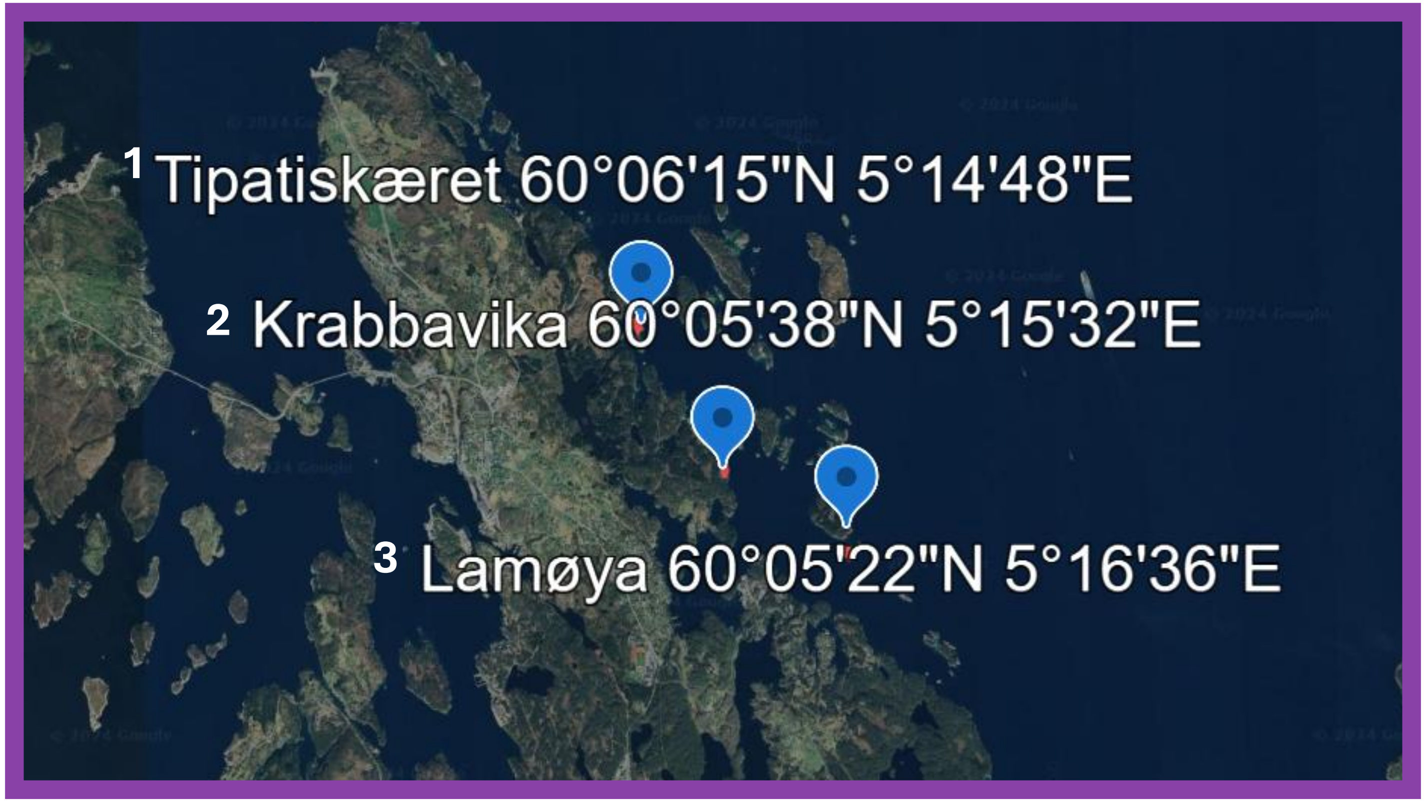


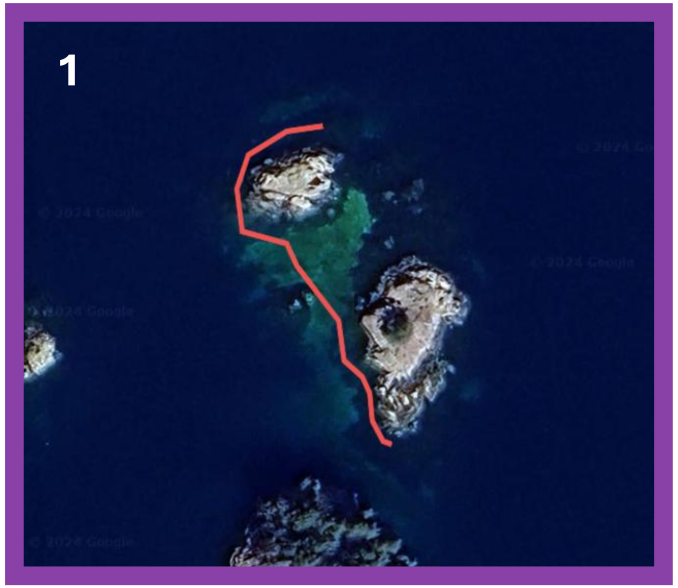


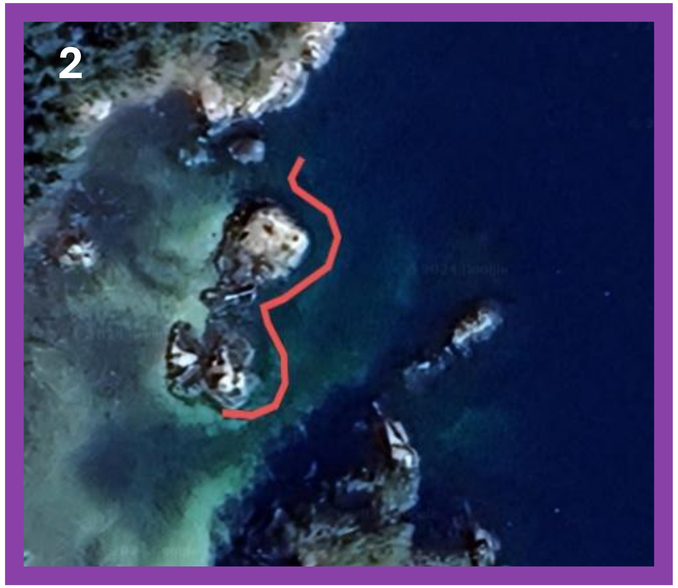


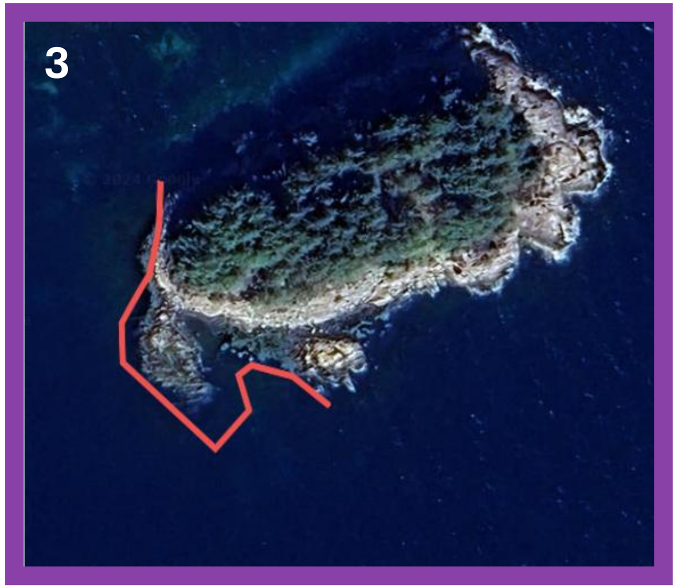

**Transect coordinates for the 3 sub-locations of the Hitra population.**

Red lines indicate artificial nests lines. 20 nests are placed along each line.

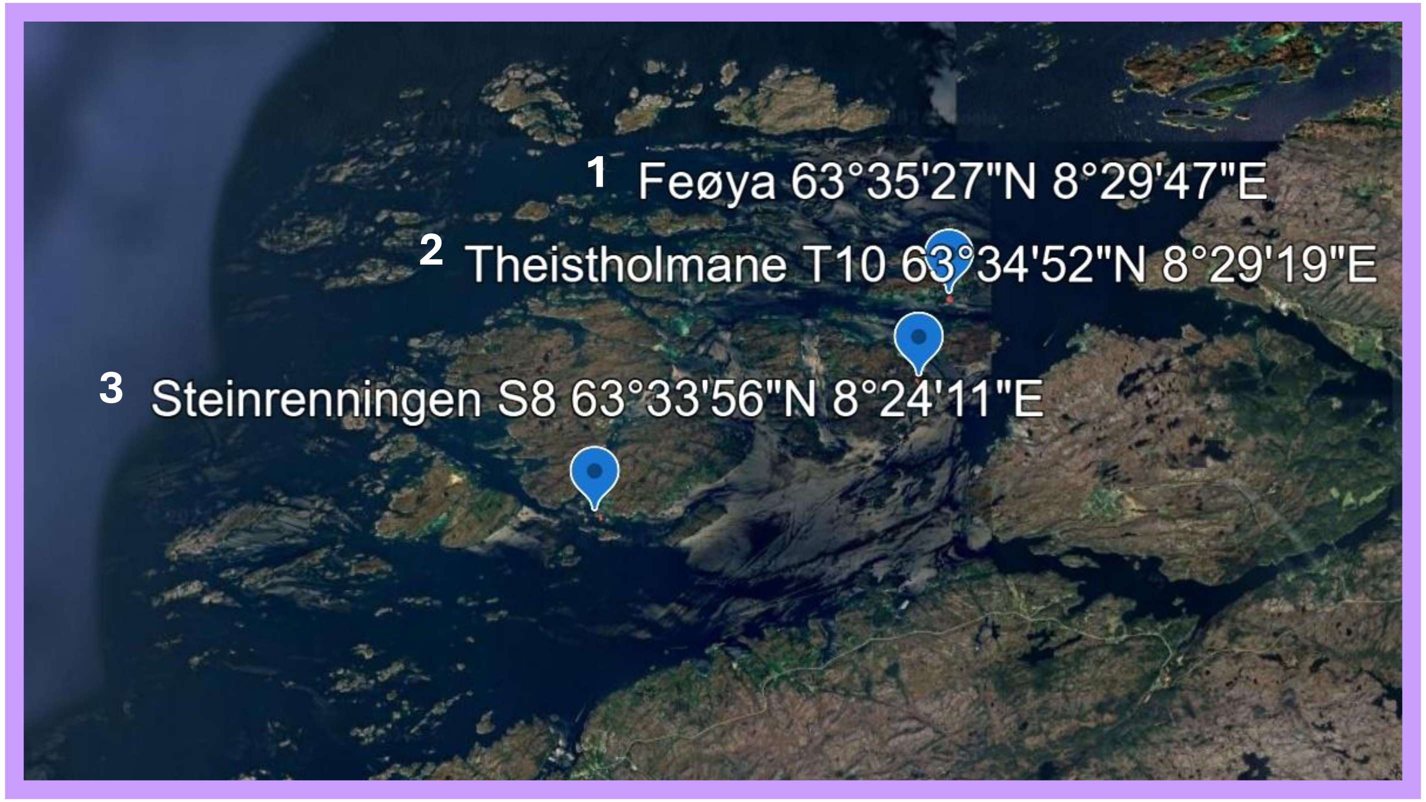


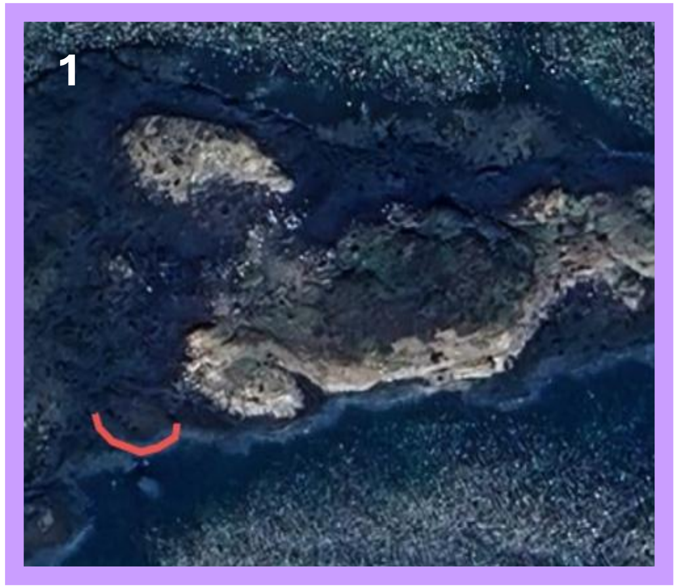


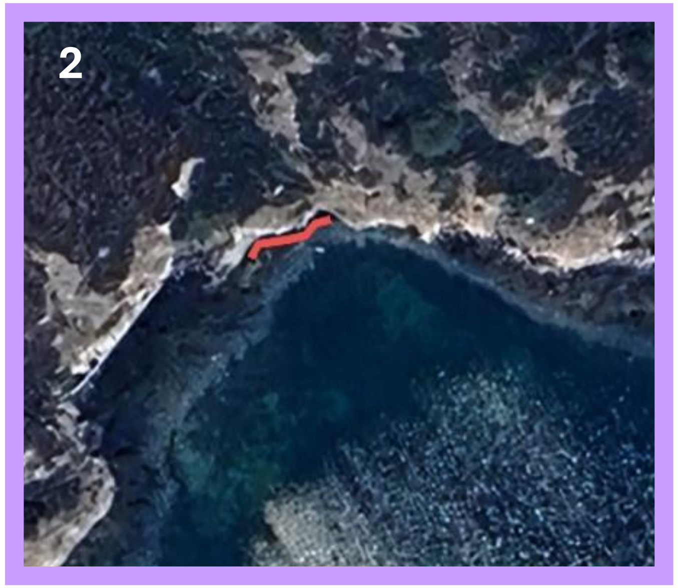


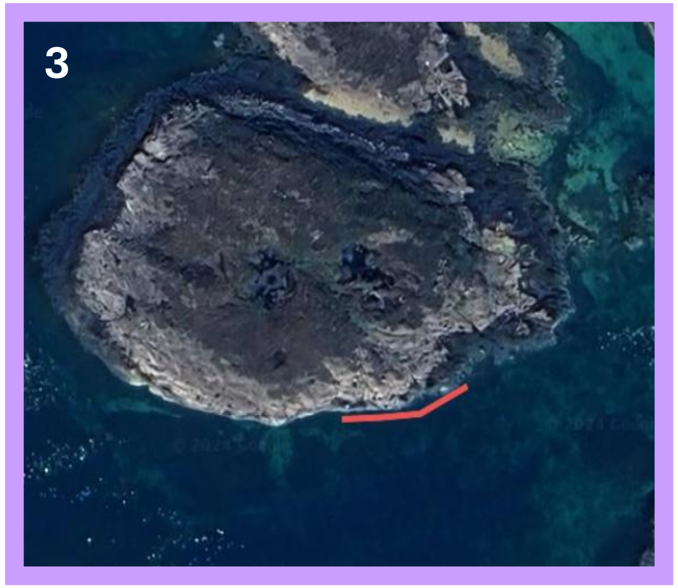

#### Transect coordinates for the 3 sub-locations of the Helligvær population.

Red lines indicate artificial nests lines. 20 nests are placed along each line.

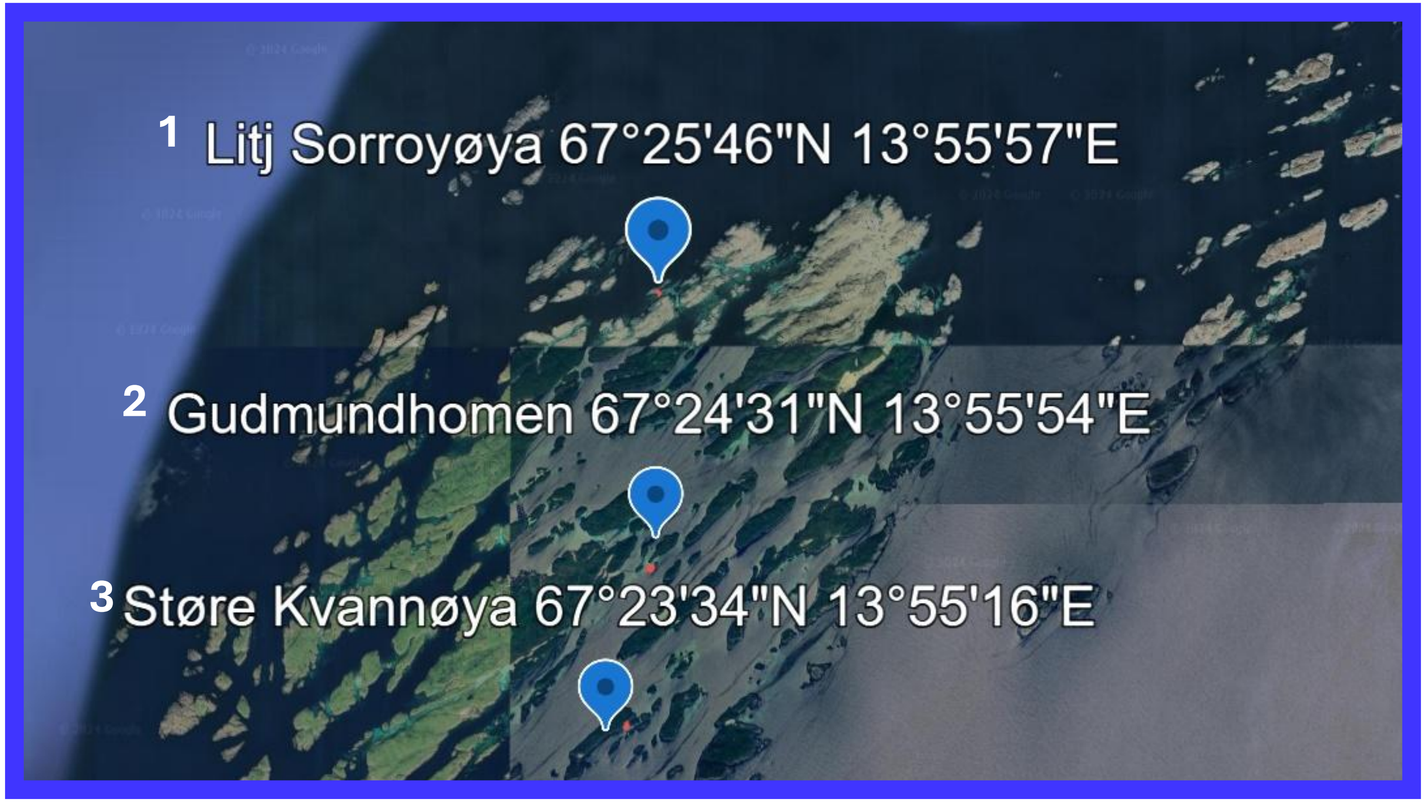


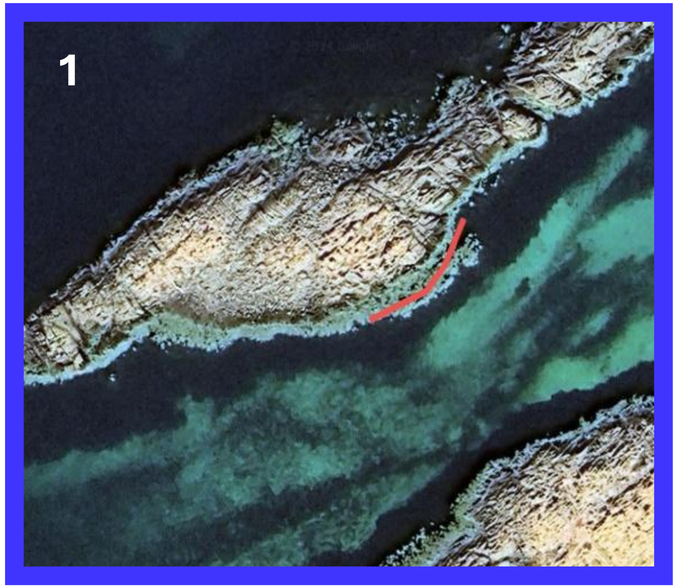


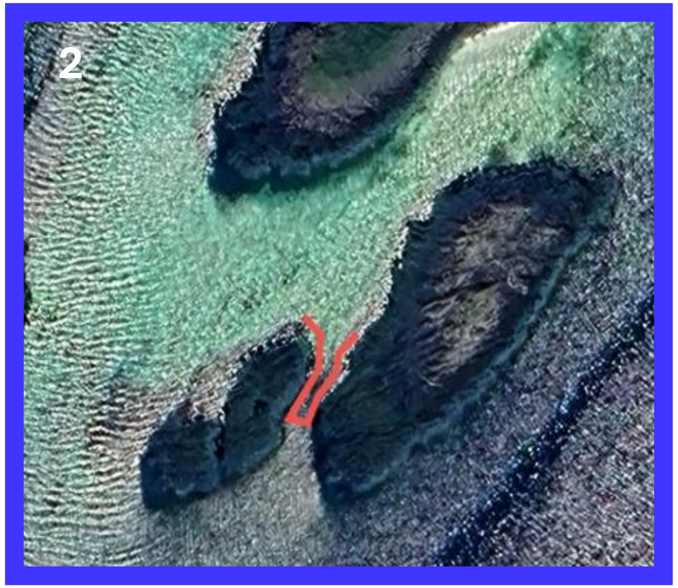


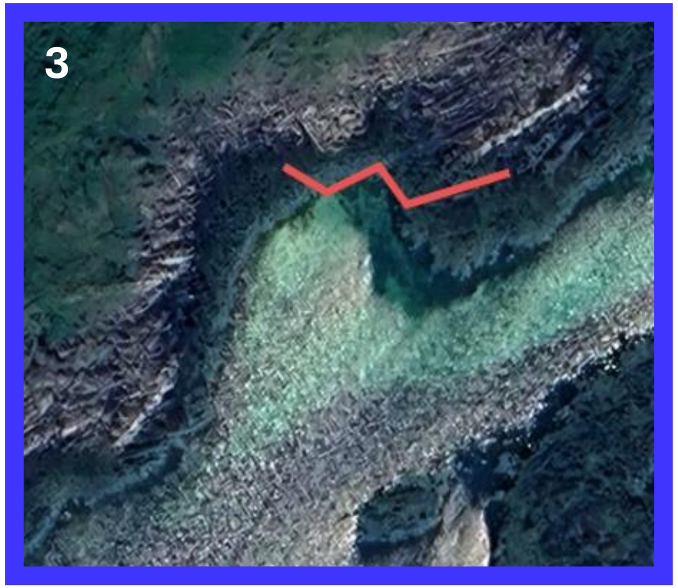

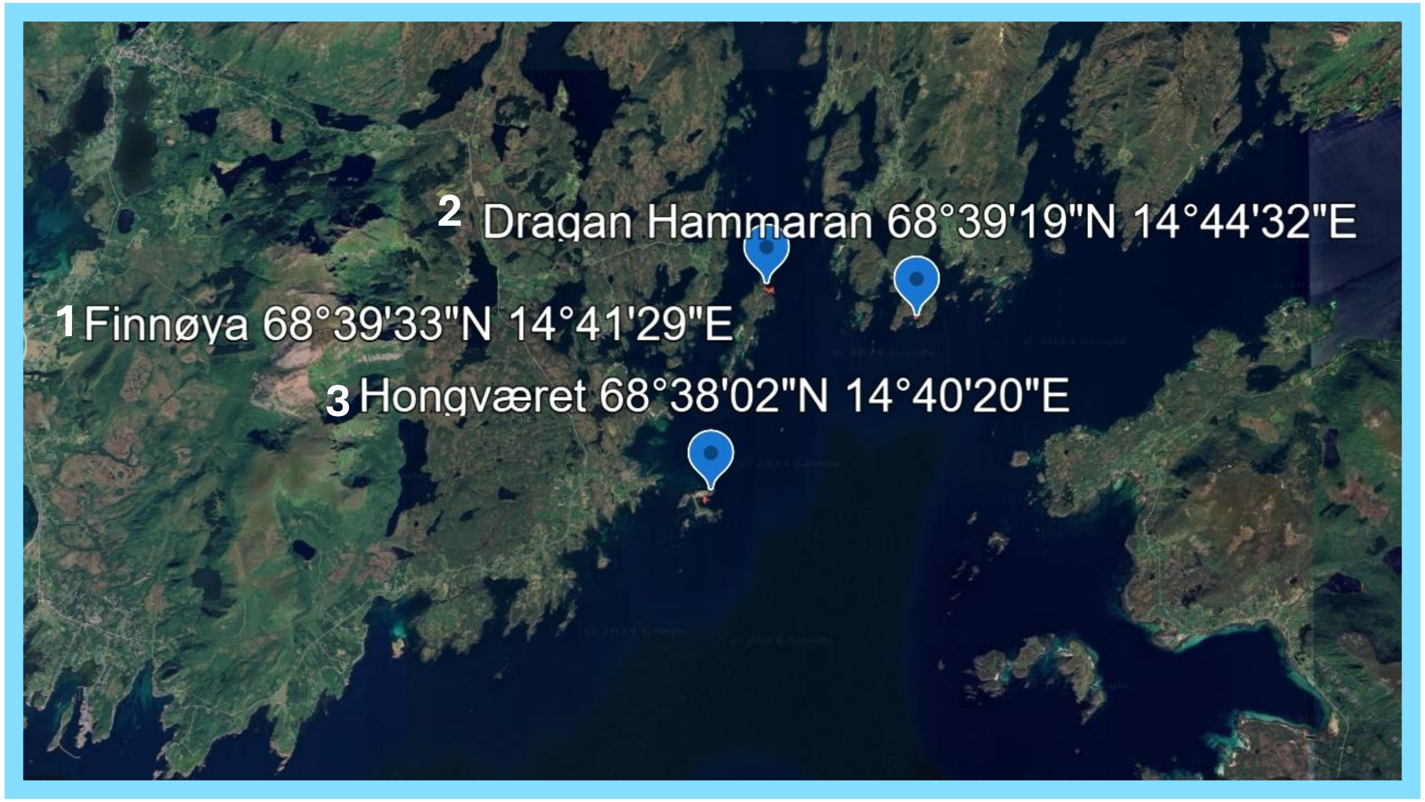


**Transect coordinates for the 3 sub-locations of the Ringstad population.**

Red lines indicate artificial nests lines. 20 nests are placed along each line.

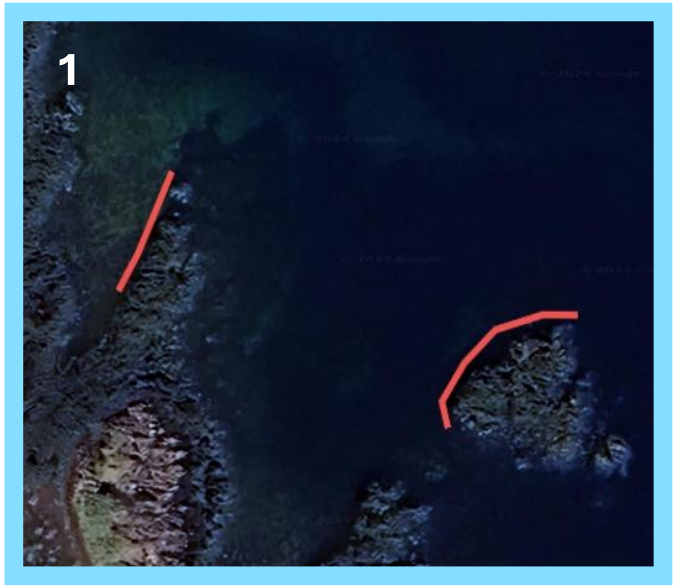


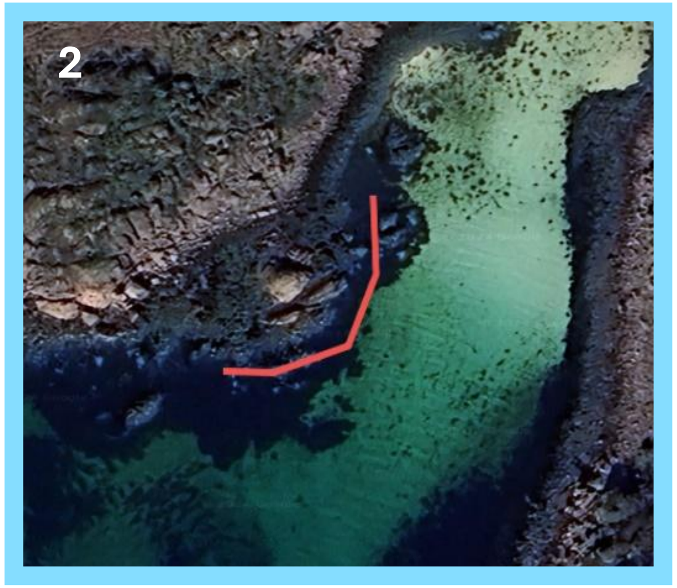


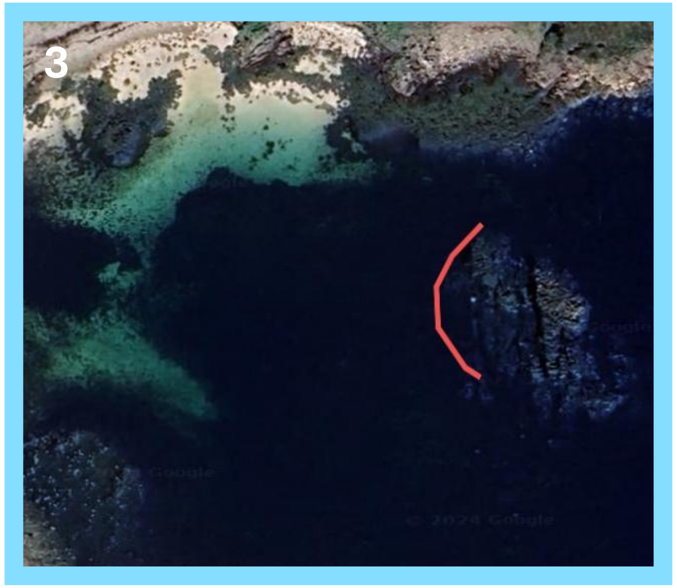
