## Supplementary tables for "Local mating competition, but not climate, drives male reproductive success across a latitudinal gradient in a nest-brooding marine fish"

**Supplementary table 1.** Model 1 of the path diagram (Figure 4.a), linear mixed effect model with reproductive success as a response, peak season data.

| **Fixed Effect** | **Estimate** | **Chi sq.** | **Df** | **p** |
| --- | --- | --- | --- | --- |
| DDF | -0.15 | 0.50 | 1 | 0.479 |
| Male length (mean) | -0.50 | 2.32 | 1 | 0.128 |
| Male condition (mean) | Excluded, collinear with DDF | | | |
| Cannibalism (mean) | 0.79 | 1.12 | 1 | 0.289 |
| Overgrowth (mean) | -0.47 | 1.95 | 1 | 0.162 |
| Nest size | 1.36 | 63.8 | 1 | <0.001 |
| Male length (rel.) | 0.30 | 9.00 | 1 | 0.003 |
| Male condition (rel.) | -0.12 | 1.47 | 1 | 0.225 |
| Cannibalism (rel.) | -0.16 | 4.26 | 1 | 0.039 |
| Overgrowth (rel.) | 0.04 | 0.22 | 1 | 0.636 |
| **Random effect** | **Variance** | |  |  |
| Sampling site | 0.08 | |  |  |
| Residual | 0.40 | |  |  |

**Supplementary table 2.** Model 1 of the path diagram (Figure 4.b), linear mixed effect model with reproductive success as a response, late season data.

| **Fixed Effect** | **Estimate** | **Chi sq.** | **Df** | **p** |
| --- | --- | --- | --- | --- |
| DDF | 0.02 | 0.01 | 1 | 0.917 |
| Male length (mean) | 0.14 | 0.26 | 1 | 0.610 |
| Male condition (mean) | 0.60 | 2.87 | 1 | 0.090 |
| Cannibalism (mean) | -0.35 | 0.70 | 1 | 0.403 |
| Overgrowth (mean) | Excluded, collinear with DDF | | | |
| Nest size | 1.29 | 50.5 | 1 | <0.001 |
| Male length (rel.) | 0.05 | 0.26 | 1 | 0.611 |
| Male condition (rel.) | -0.02 | 0.09 | 1 | 0.770 |
| Cannibalism (rel.) | -0.43 | 18.2 | 1 | <0.001 |
| Overgrowth (rel.) | -0.07 | 0.39 | 1 | 0.532 |
| **Random effect** | **Variance** | |  |  |
| Sampling site | 0.10 | |  |  |
| Residual | 0.37 | |  |  |

**Supplementary table 3.** Model 2 of the path diagram (Figure 4.a), beta regression with cannibalism index as a response, peak season data. (*) Estimates of effect size are from a linear model.

| **Fixed Effect** | **Estimate (*)** | **Chi sq.** | **Df** | **p** |
| --- | --- | --- | --- | --- |
| DDF | 0.30 | 4.23 | 1 | 0.039 |
| Male length (mean) | 0.25 | 0.55 | 1 | 0.457 |
| Male condition (mean) | Excluded, collinear with DDF | | | |
| Nest size | -0.05 | 0.03 | 1 | 0.862 |
| Male length (rel.) | -0.21 | 1.18 | 1 | 0.277 |
| Male condition (rel.) | -0.12 | 0.06 | 1 | 0.807 |
| **Random effect** | **Variance** |  |  |  |
| Sampling site | 6.3 x10^-10^ |  |  |  |

**Supplementary table 4.** Model 2 of the path diagram (Figure 4.b), beta regression with cannibalism index as a response, late season data. (*) Estimates of effect size are from a linear model.

| **Fixed Effect** | **Estimate (*)** | **Chi sq.** | **Df** | **p** |
| --- | --- | --- | --- | --- |
| DDF | 0.47 | 13.4 | 1 | <0.001 |
| Male length (mean) | 0.44 | 1.01 | 1 | 0.314 |
| Male condition (mean) | -0.28 | 1.44 | 1 | 0.230 |
| Nest size | 0.02 | 0.39 | 1 | 0.530 |
| Male length (rel.) | -0.07 | 0.14 | 1 | 0.712 |
| Male condition (rel.) | 0.05 | 0.93 | 1 | 0.334 |
| **Random effect** | **Variance** |  |  |  |
| Sampling site | 1.97 x10^-9^ |  |  |  |

**Supplementary table 5.** Model 3 of the path diagram (Figure 4.a), hurdle model (zero-inflated beta regression) with overgrowth index as a response, peak season data. (*) Estimates of effect size are from a linear model.

|  | | **Binomial (y/n)** | | | **Beta regression (>0)** | | |
| --- | --- | --- | --- | --- | --- | --- | --- |
| **Fixed Effect** | **Estimate (*)** | **Chi sq.** | **Df** | **p** | **Chi sq.** | **Df** | **p** |
| DDF | 0.23 | 1.85 | 1 | 0.173 | 5.80 | 1 | 0.016 |
| Male length (mean) | -0.75 | 4.16 | 1 | 0.041 | 7.60 | 1 | 0.006 |
| Male cond. (mean) | Excluded, collinear with DDF | | | | | | |
| Nest size | -0.57 | 0.12 | 1 | 0.726 | 10.5 | 1 | 0.001 |
| Male length (rel.) | 0.05 | 0.13 | 1 | 0.721 | 0.28 | 1 | 0.599 |
| Male condition (rel.) | -0.10 | 0.75 | 1 | 0.388 | 1.43 | 1 | 0.232 |
| **Random effect** |  | **Variance** |  |  | **Variance** |  |  |
| Sampling site |  | 4.79 |  |  | 2.2 x10^-9^ |  |  |

**Supplementary table 6.** Model 3 of the path diagram (Figure 4.b), hurdle model (zero-inflated beta regression) with overgrowth index as a response, late season data. (*) Estimates of effect size are from a linear model.

|  | | **Binomial (y/n)** | | | **Beta regression (>0)** | | |
| --- | --- | --- | --- | --- | --- | --- | --- |
| **Fixed Effect** | **Estimate (*)** | **Chi sq.** | **Df** | **p** | **Chi sq.** | **Df** | **p** |
| DDF | 0.40 | 2.20 | 1 | 0.138 | 5.47 | 1 | 0.019 |
| Male length (mean) | -0.11 | 2.69 | 1 | 0.101 | 0.78 | 1 | 0.377 |
| Male condition (mean) | 0.25 | 3.67 | 1 | 0.055 | 0.52 | 1 | 0.469 |
| Nest size | -0.79 | 1.18 | 1 | 0.278 | 12.8 | 1 | <0.001 |
| Male length (rel.) | -0.11 | 3.93 | 1 | 0.048 | 0.19 | 1 | 0.667 |
| Male condition (rel.) | -0.09 | 0.28 | 1 | 0.598 | 0.05 | 1 | 0.828 |
| **Random effect** |  | **Variance** |  |  | **Variance** |  |  |
| Sampling site |  | 0.24 |  |  | 6.7 x10^-10^ |  |  |

**Supplementary table 7.** Model 4 of the path diagram (Figure 4.a), linear generalized mixed effects model with nest quality as a binomial response, peak season data.

| **Fixed Effect** | **Estimate** | **Chi sq.** | **Df** | **p** |
| --- | --- | --- | --- | --- |
| Male length (mean) | 0.09 | 0.18 | 1 | 0.675 |
| Male condition (mean) | 0.16 | 0.59 | 1 | 0.441 |
| Male length (rel.) | 1.01 | 20.6 | 1 | <0.001 |
| Male condition (rel.) | 0.549 | 8.07 | 1 | 0.005 |
| **Random effect** | **Variance** |  |  |  |
| Sampling site | 0 |  |  |  |

**Supplementary table 8.** Model 4 of the path diagram (Figure 4.b), linear generalized mixed effects model with nest quality as a binomial response, late season data.

| **Fixed Effect** | **Estimate** | **Chi sq.** | **Df** | **p** |
| --- | --- | --- | --- | --- |
| Male length (mean) | 0.06 | 0.04 | 1 | 0.852 |
| Male condition (mean) | 0.23 | 0.24 | 1 | 0.626 |
| Male length (rel.) | 1.01 | 12.2 | 1 | 0.001 |
| Male condition (rel.) | 0.19 | 0.94 | 1 | 0.332 |
| **Random effect** | **Variance** |  |  |  |
| Sampling site | 0 |  |  |  |

**Supplementary table 9.** Model 5 of the path diagram (Figure 4.a), linear mixed effects model with male length as a response, peak season data.

| **Fixed Effect** | **Estimate** | **Chi sq.** | **Df** | **p** |
| --- | --- | --- | --- | --- |
| DDF | -0.38 | 5.80 | 1 | 0.016 |
| **Random effect** | **Variance** |  |  |  |
| Sampling site | 0.34 |  |  |  |
| Residual | 0.62 |  |  |  |

**Supplementary table 10.** Model 5 of the path diagram (Figure 4.b), linear mixed effects model with male length as a response, late season data.

| **Fixed Effect** | **Estimate** | **Chi sq.** | **Df** | **p** |
| --- | --- | --- | --- | --- |
| DDF | -0.40 | 7.83 | 1 | 0.005 |
| **Random effect** | **Variance** |  |  |  |
| Sampling site | 0.30 |  |  |  |
| Residual | 0.63 |  |  |  |

**Supplementary table 11.** Model 5 of the path diagram (Figure 4.a), linear mixed effects model with male condition as a response, peak season data.

| **Fixed Effect** | **Estimate** | **Chi sq.** | **Df** | **p** |
| --- | --- | --- | --- | --- |
| DDF | 0.59 | 76.1 | 1 | <0.001 |
| **Random effect** | **Variance** |  |  |  |
| Sampling site | 0.01 |  |  |  |
| Residual | 0.64 |  |  |  |

**Supplementary table 12.** Model 5 of the path diagram (Figure 4.b), linear mixed effects model with male condition as a response, late season data.

| **Fixed Effect** | **Estimate** | **Chi sq.** | **Df** | **p** |
| --- | --- | --- | --- | --- |
| DDF | 0.53 | 46.4 | 1 | <0.001 |
| **Random effect** | **Variance** |  |  |  |
| Sampling site | 0.0 |  |  |  |
| Residual | 0.73 |  |  |  |
