## Supplementary file F2 for "Local mating competition, but not climate, drives male reproductive success across a latitudinal gradient in a nest-brooding marine fish"

### Part 1. Nest picture analysis protocol



1. Check field form for exclusion criterions:
   i. reported presence/absence of male -> if absent, caution for signs of abandoned brood (abandonned broods are discarded. Identified by mushy looking egg masses, whitish coloration of eggs)

   IF the nest looks abandoned, take note in the comment column and only measure total brood area and area of one square. The rationale is that any overgrowth or eating of eggs that might have happened after the departure of the male is not comparable to overgrowth or cannibalism in a nest with attending males.

   i. reported live eggs? -> if noted as NOT live eggs, discard from analysis
   ii. reported hatching of eggs in the field-> if noted as hatched in the field, discard from analysis, unless a good picture pre-hatching from the field is available.

2. Open nest image in imageJ. If needed, adjust brightness and contrast
3. Define and measure brood area (see slide 2 ).
4. Define and measure the area of 1 square on the pattern printed on the acetate sheet (see slide 3).
5. Define and measure up to 10 largest empty areas within the brood area, if any (assumed cannibalism, see slide 4)
6. Define and measure up to 10 largest overgrown areas within the brood, if any (see slide 5).
7. Measure egg density: create a small square shape and measure its area. Move the shape across the image five times, and count egg number within the shape each time. The shape must be placed within the brood, outside of the already defined cannibalized or overgrown areas (see slide 6), and as randomly as possible.
8. If eggs are very advanced in development, eyed and transparent, take note of it in the comment column. Missing eggs could be due to hatching and we can exclude these broods from the cannibalism data.

### Nest picture analysis protocol



3. Defining and measuring brood area

-using the freehand selection tool, follow the outline of the brood (in yellow on the pictures)

-follow outer edges, even if presenting irregular shape. The outer edge is assumed to represent the egg laying pattern (not caused by cannibalism).

-Press Ctrl+T to add the shape to the ROI manager. Press Ctrl+M to measure the area in pixels

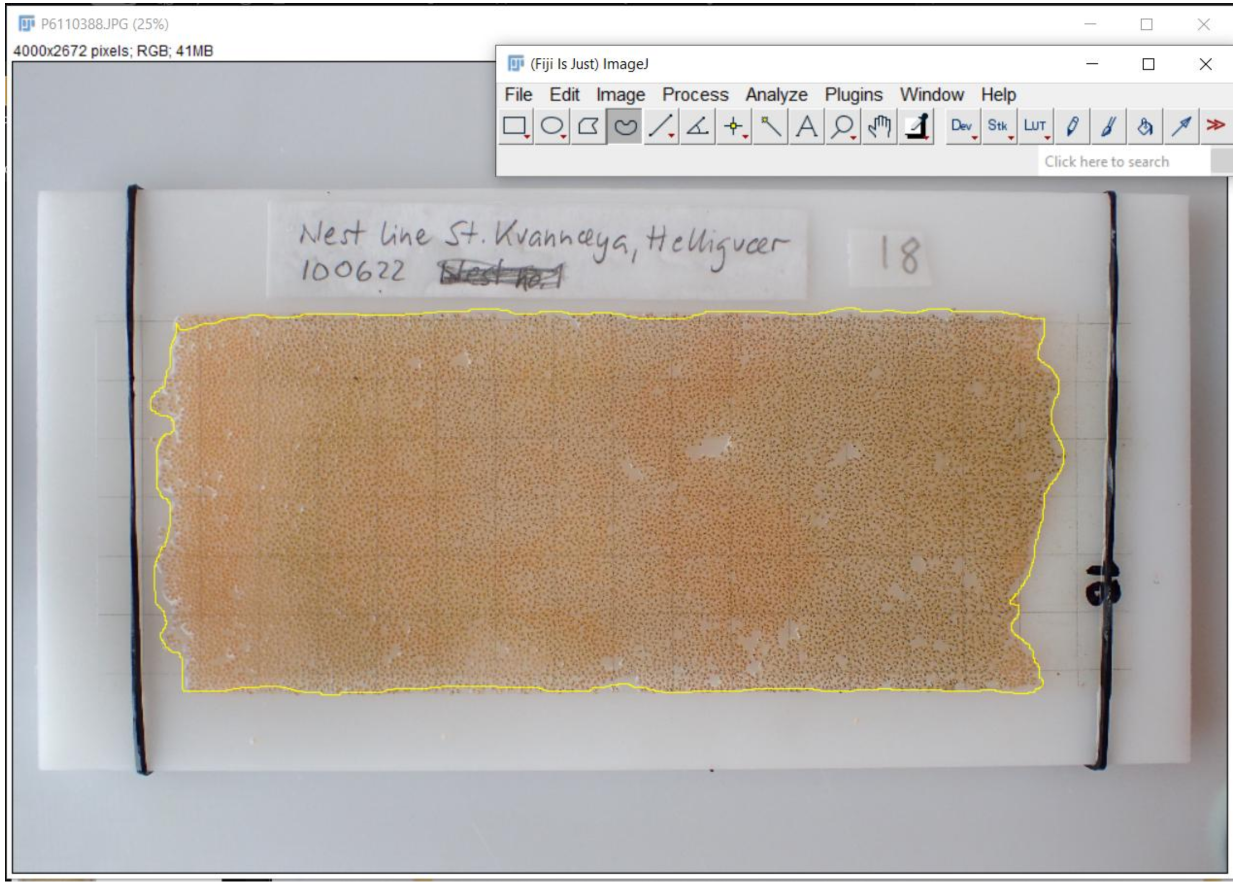


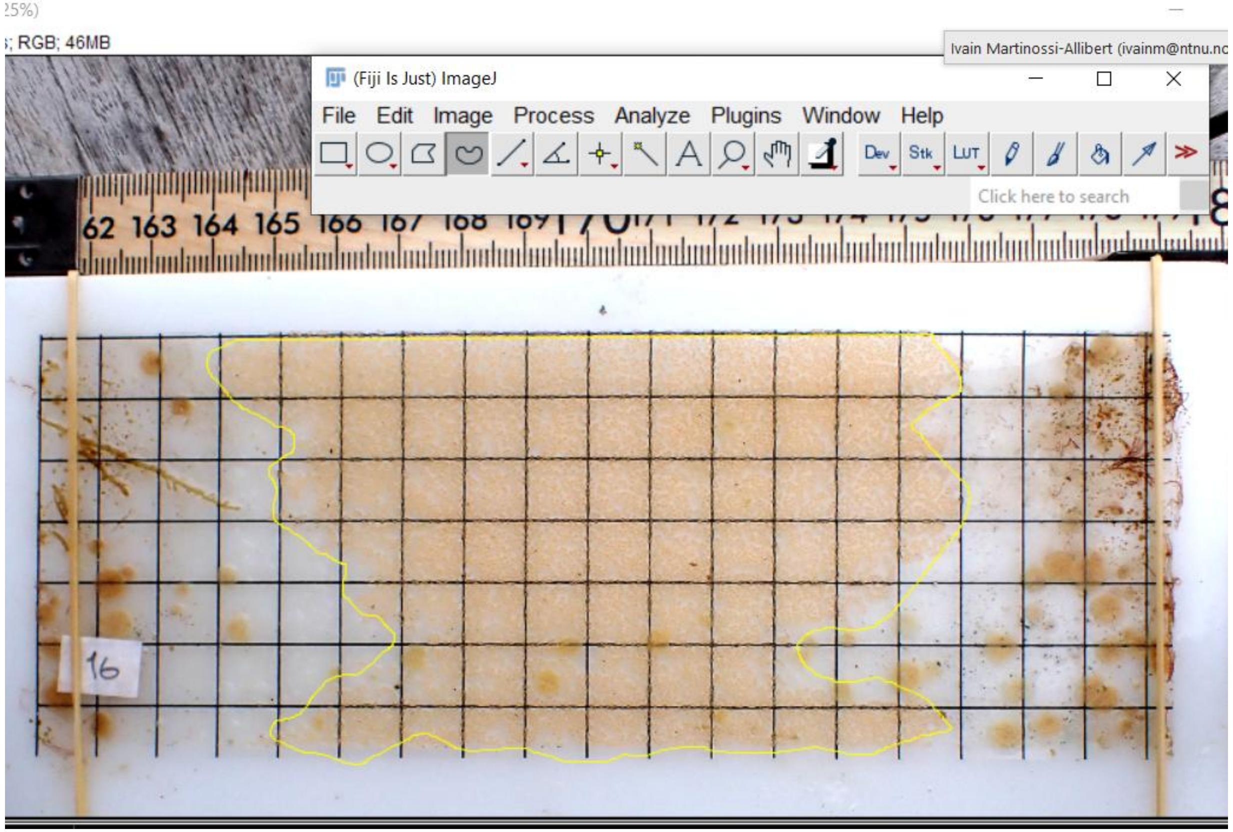

### Nest picture analysis protocol



4. Define and measure area of 1 square
-Use the square shape selection tool
-Press Ctrl+T to add the shape to the ROI manager. Press Ctrl+M to measure the area in pixels

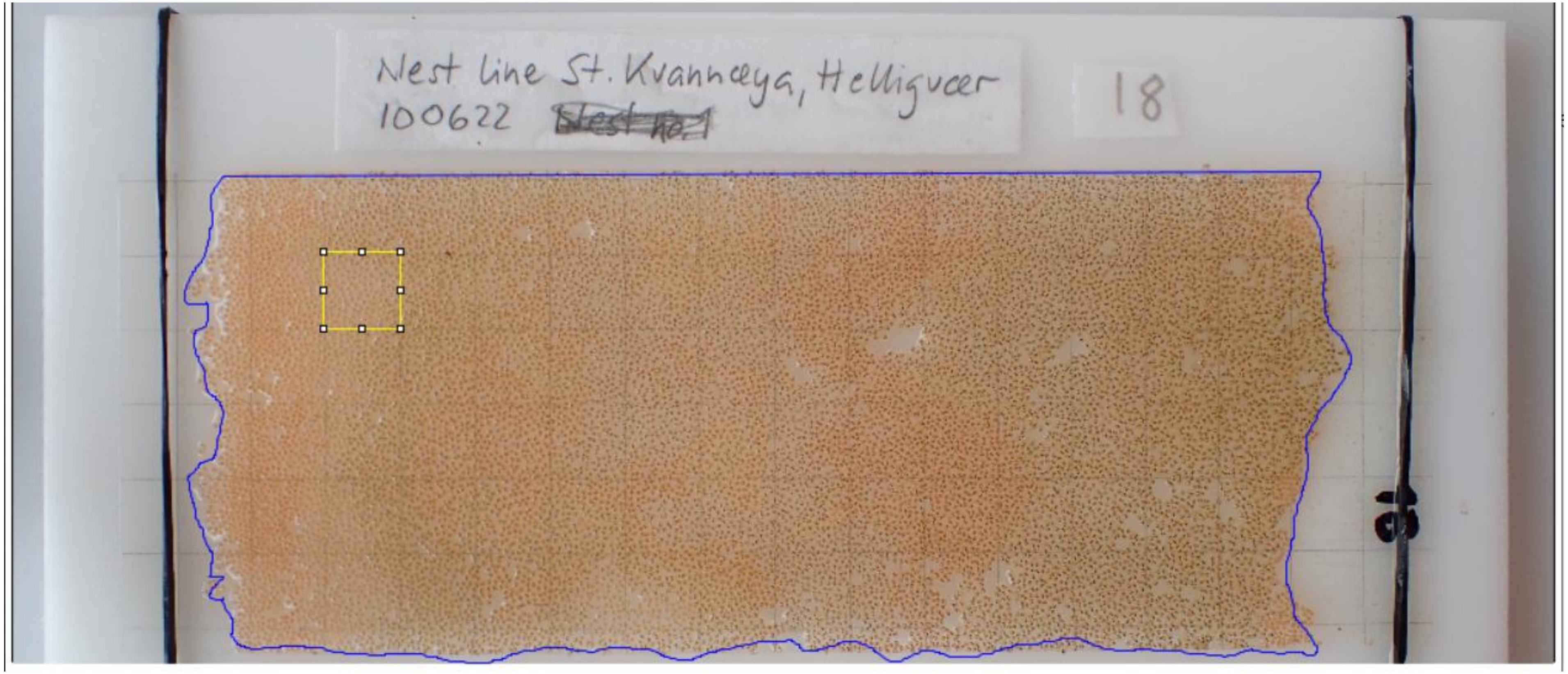

### Nest picture analysis protocol



5. Define and measure 10 largest empty areas within the brood (assumed cannibalism)
-Use free hand selection tool
-After each selection, press Ctrl+T to add the shape to the ROI manager
-Once all desired empty areas are selected, select them in the ROI manager, right click, choose “OR(Combine)”
-Then press Ctrl+M to measure total surface of combined areas.

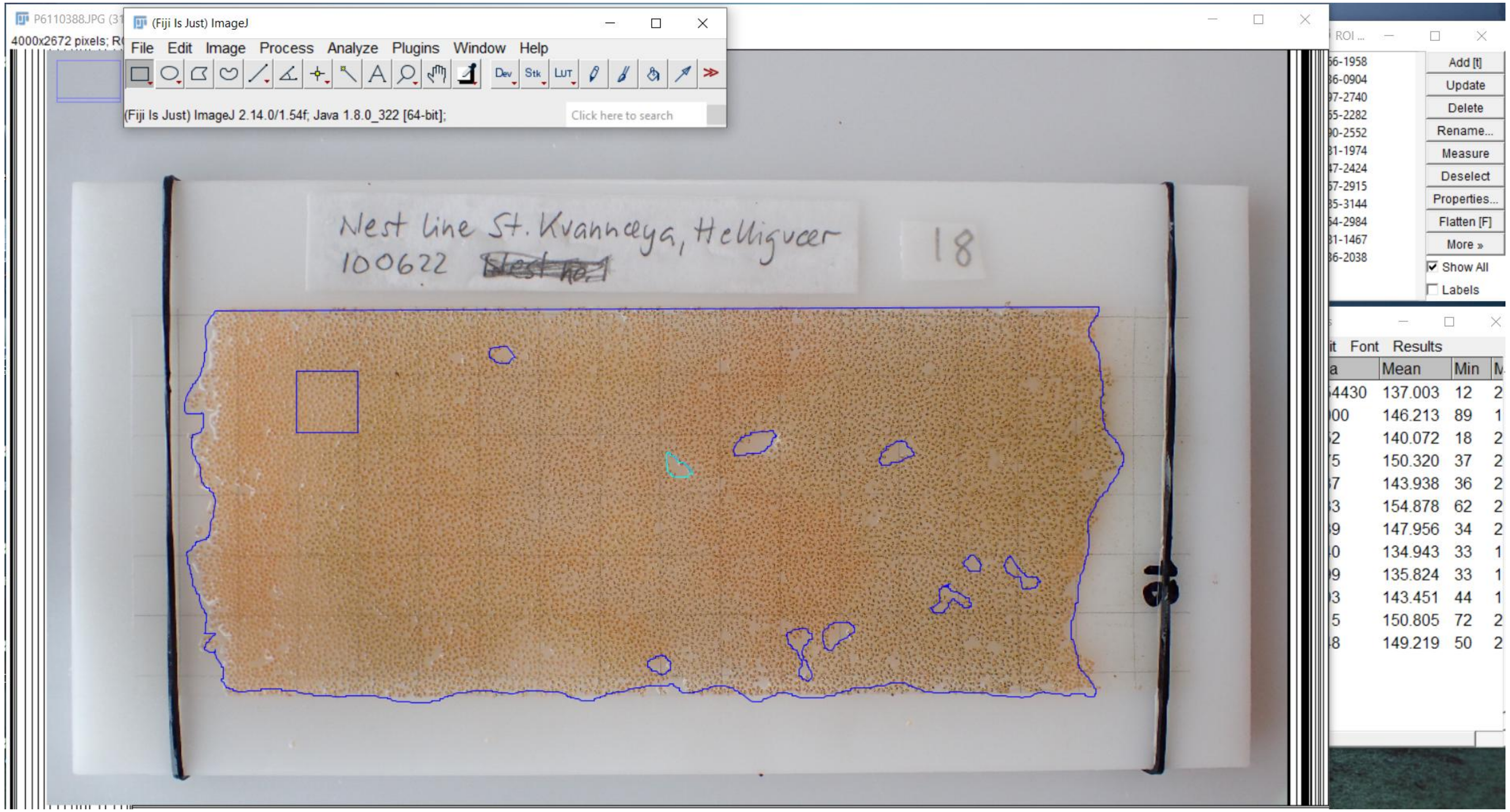

### Nest picture analysis protocol



6. Define and measure overgrown areas within the brood (up to 10 areas)
-Use free hand selection tool
-After each selection, press Ctrl+T and Ctrl+M

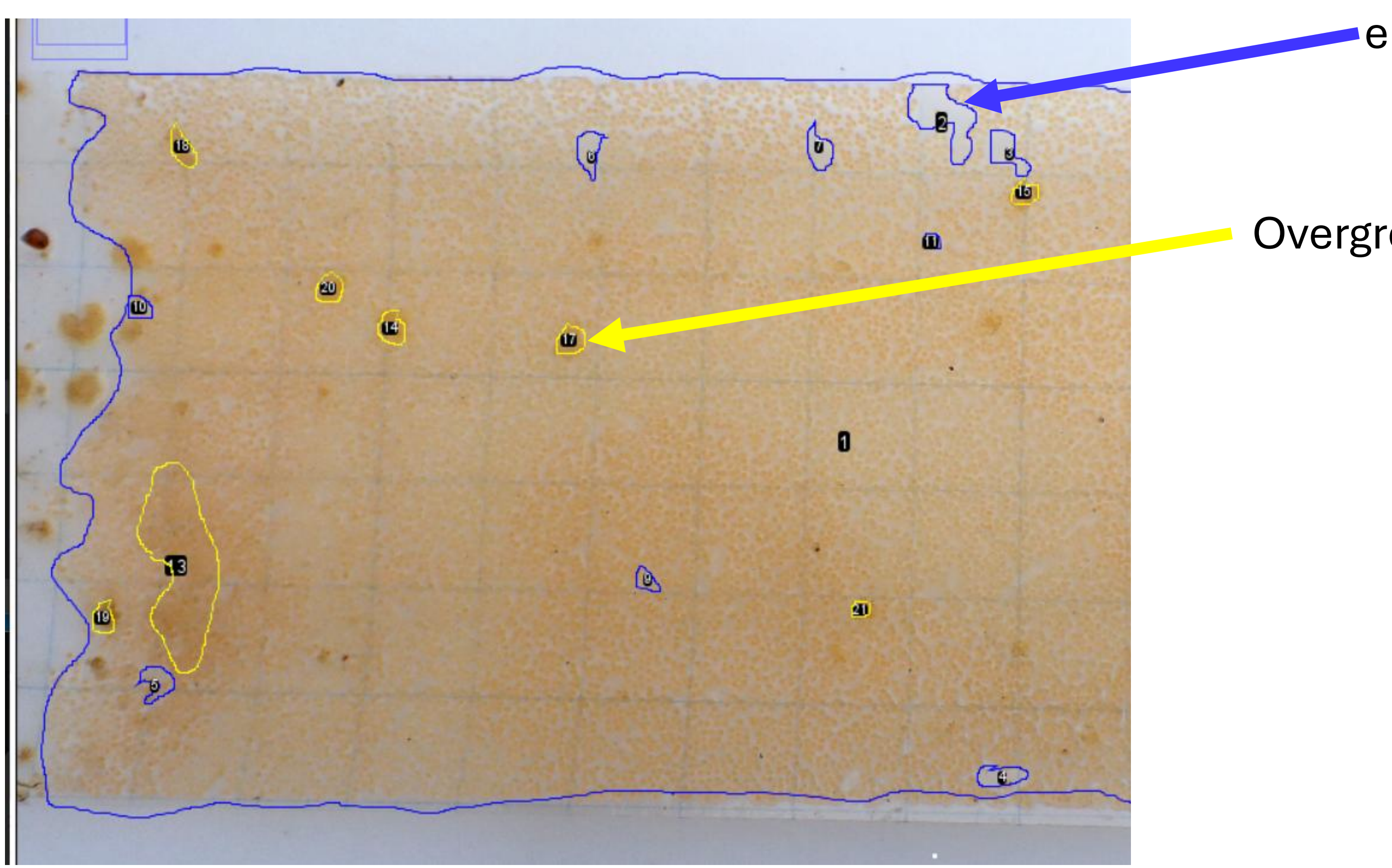

### Nest picture analysis protocol



7. Measure egg density within the brood
-Create small square shape with the square selection tool
-Measure area of shape with Ctrl+M
-Count egg number within shape (all eggs that intersect with the edge of the shape are counted. Counting is done by placing a white dot over a counted egg.
-Move the shape around and repeat counting four additional times. The shape can be placed anywhere within the brood area, but outside of the cannibalized or overgrown areas previously defined.

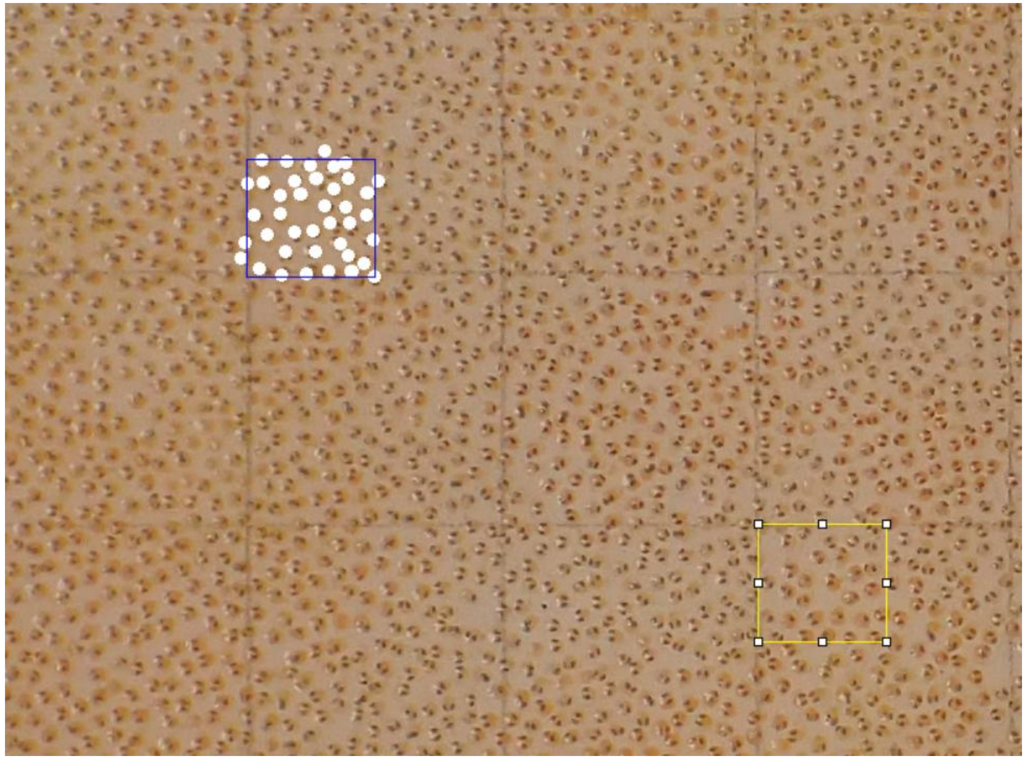

### Nest picture analysis protocol



#### Example of picture analysis

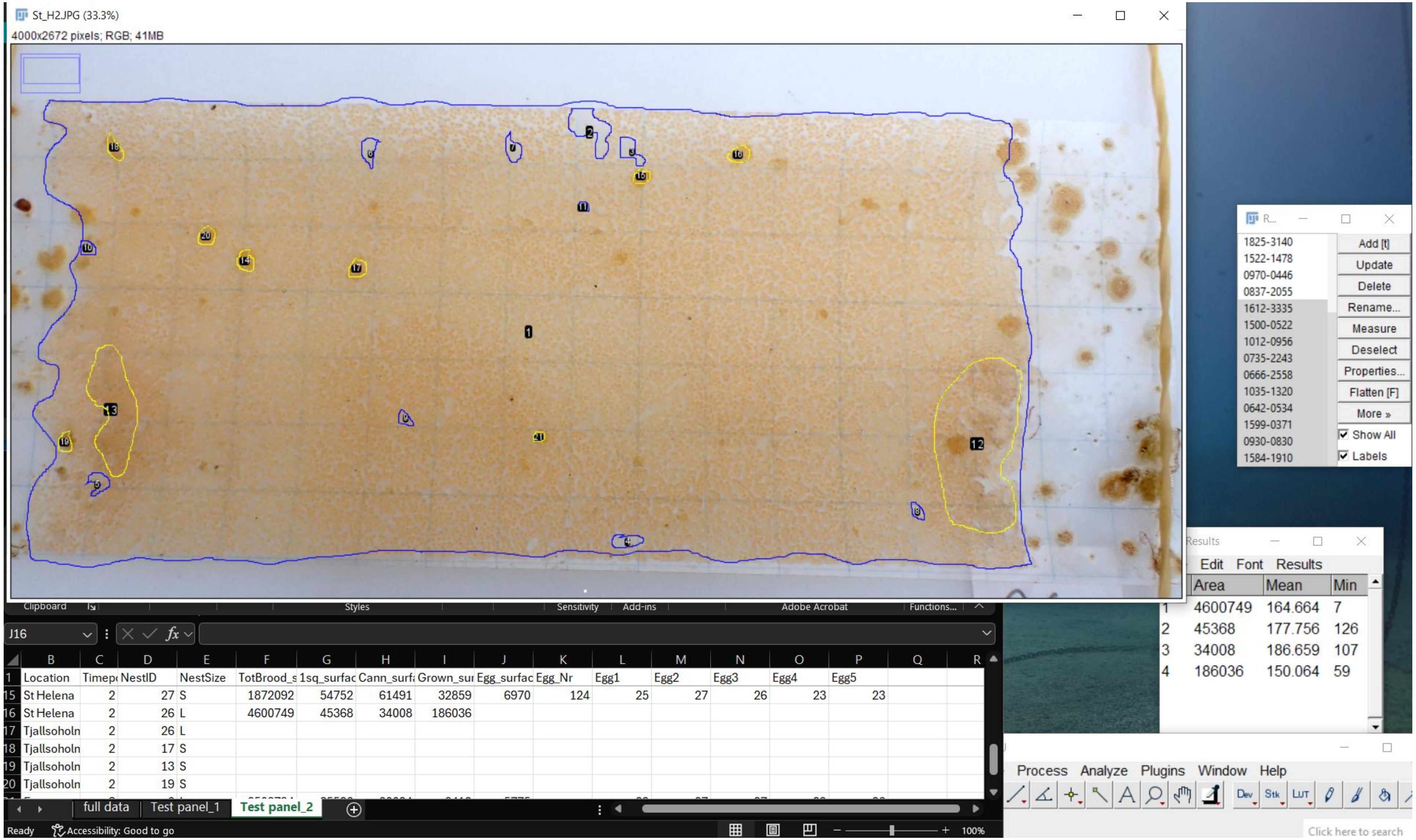


Total brood area and cannibalized areas in blue, overgrown areas in yellow.

### Part 2. Repeatability of nest picture analysis



To test for repeatability of the nest picture analysis, 20 images from different locations and time points were chosen to form a test panel that was analysed twice. The analysis was done by the same person, the 20 images were analysed consecutively, the two replicate analysis performed a few days apart.

In units of squares (grid), the surface covered by the brood once the cannibalized areas and overgrown areas are subtracted

The number of eggs on the image estimated from the surface covered by eggs and 5 small areas where eggs are counted manually

The proportion of brood area that appears cannibalised.

The proportion of brood area that is overgrown. The generally low proportion (<10%) could explain the low repeatability

The total area covered by the brood, including spaces empty of eggs (cannibalism, overgrowth….)

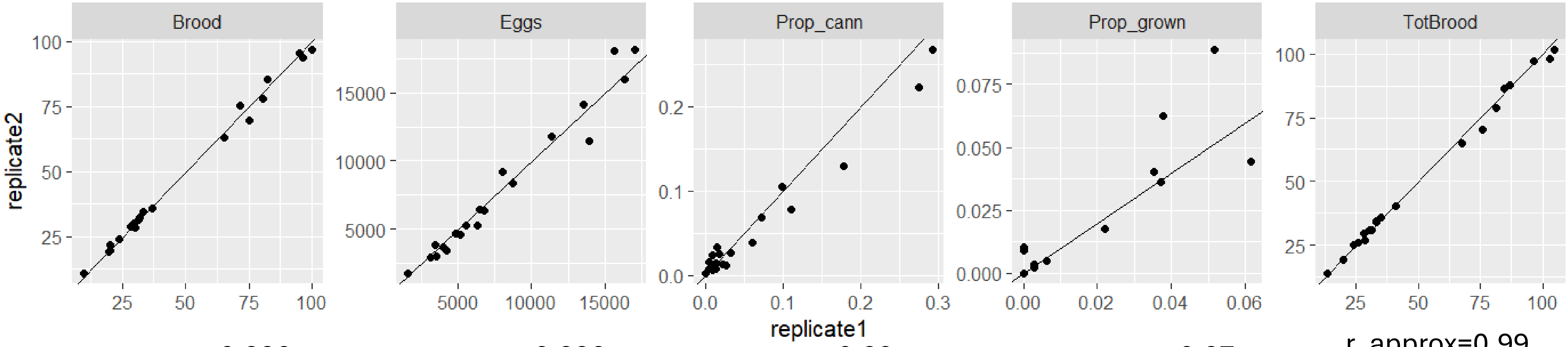


r_approx=0.996

r_approx=0.996

r_approx=0.80

r_approx=0.87

r_approx=0.99

repeatability was estimated by running a linear model and an ANOVA. For cannibalized and overgrown, the repeatability was calculated for the absolute surface in unit of grid square, not for the proportion as shown on the plot.

### Part 3. Egg diameter measurement protocol

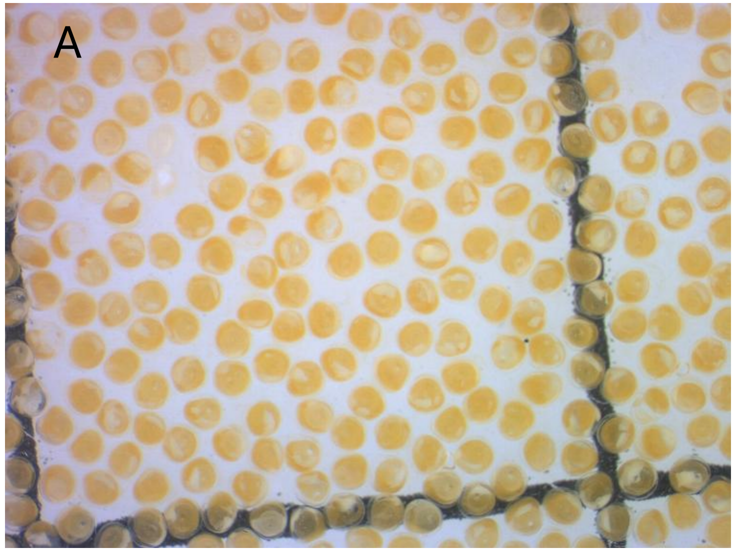


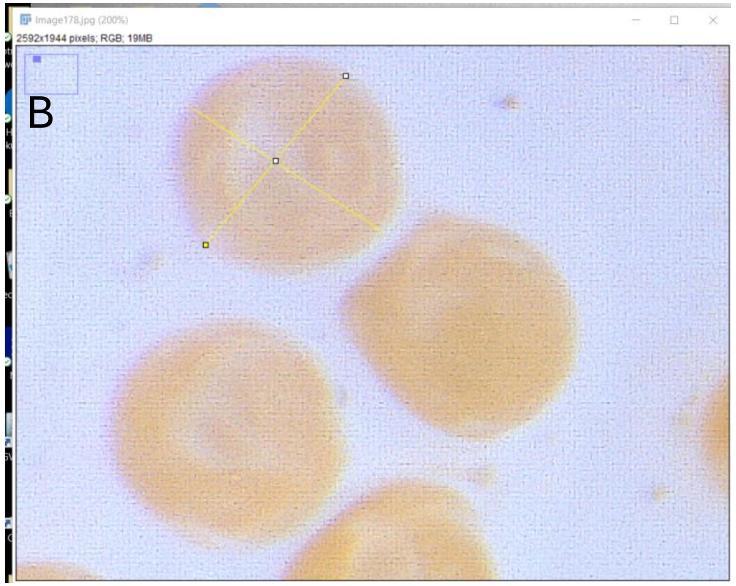


1. Picture taking
   i. When starting a session, photograph the microscope slide with engraved scale first.
   ii. Extract the acetate sheet carrying the eggs from the falcon tube, let it drain of alcohol in a petri dish for a few seconds, and lay it flat under the stereomicroscope. The acetate sheet will curl up and need to be held down flat with weights (blocks of Plexiglas where used in this case).
   iii. Take the picture (example of full picture in A)

2. Egg measurement
   1. Open the nest picture in image J. There are typically many eggs to choose from (Panel A). Zoom in on a randomly chosen location (Panel B)
   2. Measure two orthogonal diameters on a randomly picked egg. Avoid obviously damaged eggs.
   3. Measure two diameters on 5 eggs per picture.
